# Resistance evolution in a single species reorganizes community responses across a resource-antibiotic landscape

**DOI:** 10.64898/2026.11.20.768032

**Authors:** Inga-Katariina Aapalampi, Anna M Bischofberger, Dovydas Kičiatovas, Mikko Kivikoski, Ville Mustonen, Teppo Hiltunen, Johannes Cairns

## Abstract

Rapid evolution in one species can restructure ecological communities, but how its effects vary across environmental conditions remains poorly understood. Here we mapped the consequences of antibiotic resistance evolution in *Aeromonas caviae* across 64 combinations of resource availability and antibiotic stress in three- and 16-species bacterial communities. Resistance evolution changed where community-state boundaries were located, with boundaries lost or gained depending on resource availability. The same evolutionary change increased diversity in some environments but decreased it in others. Species success was partly explained by growth traits at zero or low antibiotic concentrations and by intrinsic resistance at inhibitory concentrations. Resistance evolution reversed dominance between the two dominant species and changed which traits mattered. It also altered other species’ responses below inhibitory concentrations, mostly between closely related species. Our findings show that the consequences of evolution depend on environmental context but can be partly explained by species traits.

## Introduction

Community composition depends on environmental conditions and interactions between species. Because these interactions can change across environments, the same species can assemble into different communities under different conditions. Environmental change can shift community composition and, in some systems, produce transitions between alternative states^1,2^. Ecological and evolutionary processes can also interact over similar timescales^3,4^: environmental change alters selection pressures, while evolution can modify species growth, interactions and community dynamics. Predicting community responses therefore requires understanding how evolutionary change affects the relationship between environmental conditions and community composition.

Microbial communities are a tractable system for this problem, and predicting how they respond to disturbance matters for human health, for agriculture and for environmental management^5^. We use disturbance in its standard sense, as a causal event that produces a discrete change in the physical or chemical environment^6^, and distinguish it from stress, the external constraint that conditions place on the rate of growth^7^. Antibiotic exposure is both: applied continuously it is a press disturbance that constrains growth across the community, and above inhibitory concentrations it also reduces biomass.

Environments differ in how much stress they impose, and community assembly processes shift along stress gradients. As conditions become harsher, the sign and strength of interactions between species can change^8^, and community composition can become more strongly determined by the abiotic environment than by interactions between species^5^. However, abiotic conditions and competition are not cleanly separable^9^. Where a community is located along a stress gradient should therefore shape how much evolution in any one species matters. Antibiotics affect not only their target species but entire microbial communities, altering community composition and interactions among community members^10,11^. These disturbances can destabilize microbiomes and shift communities away from their original state, weakening the community’s ability to suppress opportunistic or resistant pathogens^10–12^. Microbiome responses to antibiotic disturbance may therefore involve changes in both community stability and species interactions^12,13^. Since other microbes influence the antibiotic response of pathogens, the susceptibility and evolutionary responses of pathogens can depend strongly on the surrounding microbial community^13^.

The effects of antibiotic resistance depend on environmental and community context. The same resistance mutation can have different fitness effects across environments and communities^14–16^, sometimes reducing fitness in one host while increasing it in another^15^. These differences arise partly because resistant and susceptible cells compete for shared resources. Resource availability influences bacterial growth and competition, including competition between resistant and susceptible strains under different patterns of resource supply^14^. It also affects microbial interactions, diversity and community stability^17^.

Rapid evolution in one species can also affect the wider community^18^. Phenotypic diversity within an evolving species can change community structure by facilitating or inhibiting other taxa. Evolution can therefore alter the performance of other species and their interactions^4,18^, changing individual abundances and potentially shifting the state of the entire community^13,14^. The magnitude of these effects may depend on the phenotypic diversity that evolves within the focal species.

It remains unclear how the ecological effects of evolution in one species vary across environmental conditions^4,18,19^. Most studies examine evolutionary effects in only one or a few environments. Consequently, we do not know whether evolution produces similar community changes across environmental gradients or alters the relationship between environmental conditions and community structure. It is also unknown whether evolution in one species changes which species traits are associated with competitive success along such gradients. Resolving this question is important for predicting community responses to antibiotic disturbance and the ecological consequences of resistance evolution.

We addressed this question using a synthetic bacterial community whose properties make it well suited to it. Its response to antibiotic disturbance is repeatable^20^, it responds to resource availability with distinct community states^21^, and it contains a focal species, *A. caviae*, whose resistance evolution is known to matter at the community level^19,22^. *A. caviae* is one of the dominant species and is ancestrally moderately susceptible to streptomycin, whereas *C. koseri* is intrinsically resistant^19,22^. Because the balance between these two species shifts with streptomycin^19,22^, evolution in a single species can plausibly reshape the whole community. This makes the system suitable for asking how evolution in one species changes community responses across environmental gradients, and how these responses are linked to species traits and interactions.

We tested whether resistance evolution in *A. caviae* changed how bacterial communities respond to the joint effects of resource availability and antibiotic stress. We compared three- and 16-species communities containing either ancestral or resistance-evolved *A. caviae* across a factorial landscape of carbon and streptomycin concentrations (Fig. 1A,B). We examined community composition, diversity and the spatial organization of community states across the landscape, and whether these changes were associated with the environmental sensitivity of the focal species (Fig. 1C). Complementary monoculture assays measured growth traits across the same landscape (Fig. 1D). Comparing these traits and each species’ intrinsic resistance with species success in communities allowed us to test how community responses were linked to species traits and interactions, and whether resistance evolution changed these links.

**Figure 1.**
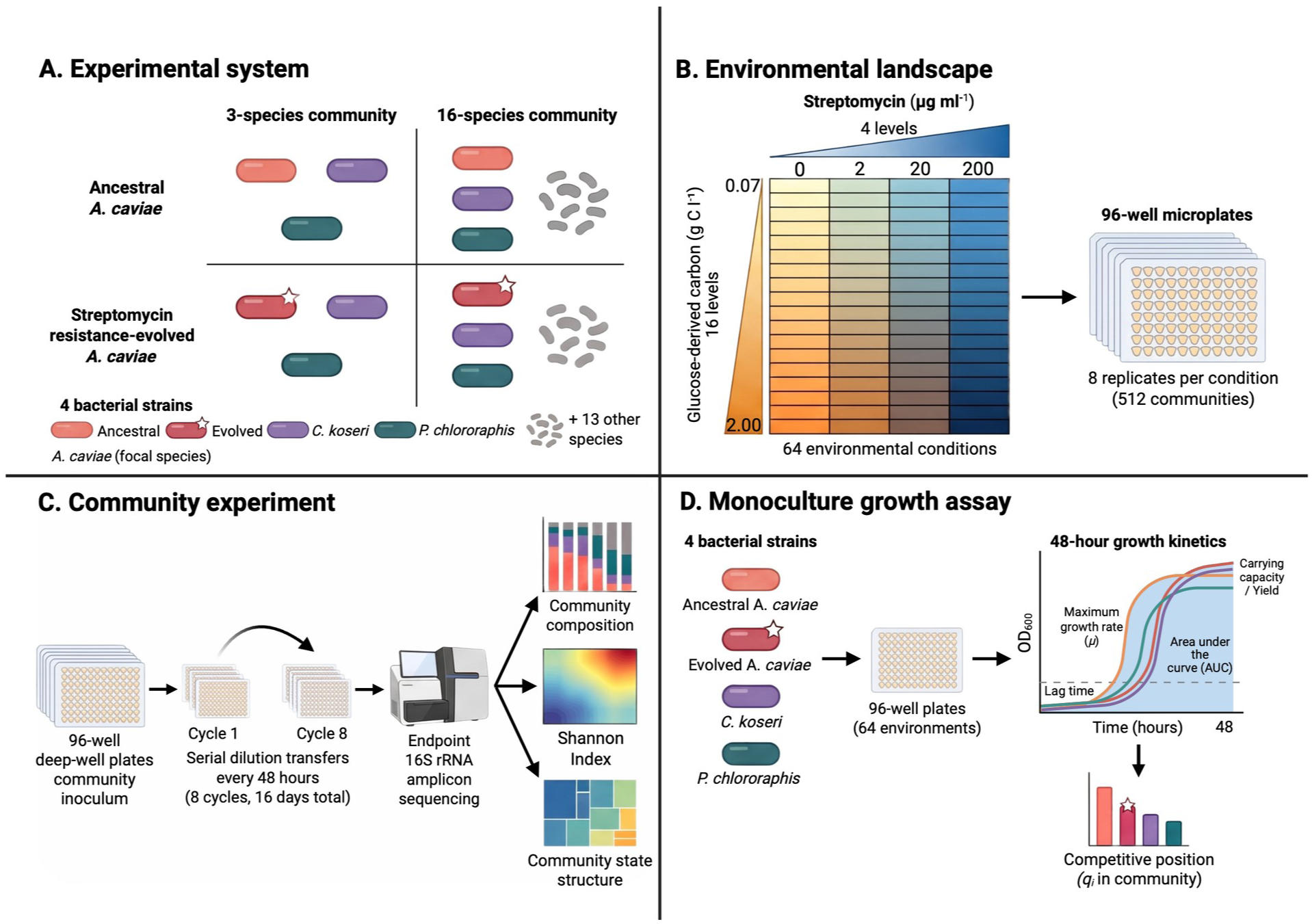
Experimental design. **A**, The four community types. Three- and 16-species communities were assembled containing either ancestral or streptomycin-resistance-evolved *Aeromonas caviae* HAMBI 1972 as the focal species (starred when resistance-evolved), together with *Citrobacter koseri* HAMBI 1287 and *Pseudomonas chlororaphis* HAMBI 1977 and, in the 16-species communities, 13 further species. The two histories of a community differed only in the *A. caviae* strain. **B**, The environmental landscape: 16 glucose-derived carbon concentrations (0.07-2.00 g C L^-1^) fully crossed with four streptomycin concentrations (0, 2, 20 and 200 µg ml^-1^), giving 64 conditions, each with eight independent replicate populations per community type. **C**, The community experiment. Populations were propagated in 96-well deep-well plates through eight 48-h serial transfers (16 days). Endpoint composition was determined by 16S rRNA gene amplicon sequencing and used to derive species relative abundance, Shannon Index and the segmented community-state structure of each landscape. **D**, The monoculture growth assay. Ancestral and resistance-evolved *A. caviae*, *C. koseri* and *P. chlororaphis* were grown alone across the same 64 environments, and optical density at 600 nm followed over 48 h was used to estimate maximum growth rate, lag time, yield and area under the curve. These traits were related to each species’ endpoint competitive position *q_i_* in the three-species communities. Created in BioRender.

We expected resistance evolution in *A. caviae* to weaken the dominance of *C. koseri* under streptomycin, and resource availability to modify this effect through its influence on *A. caviae* growth^21^. Resistance evolution could also change community diversity and state structure, altering dominance, coexistence or shifts between community states. Finally, we expected the traits underlying competitive success to vary across the landscape.

## Results

### Resistance evolution reorganizes community-state landscape topology

We first investigated whether evolution of a single community member changed the large-scale organization of endpoint (day 16, after eight 48-h serial transfers) community composition across the resource-antibiotic landscape (16 glucose-derived carbon concentrations crossed with four streptomycin concentrations) (Fig. 2A; Supplementary Figs. S1-S4; Supplementary Table S1). Landscape statistics below are means across the eight replicate landscapes of each focal-species history. Replacing ancestral *A. caviae* HAMBI 1972 with its resistance-evolved derivative changed where community-state boundaries were located. The configuration of state boundaries differed between focal-species histories in both the 16-species community (exact PERMANOVA on replicate boundary maps, R^2^=0.31, *P*=0.00016; Fig. 2B) and the three-species community (R^2^=0.32, *P*=0.00016), and comparing whole partitions gave the same result (Supplementary Table S1). Replicate landscapes with the same history shared more of their boundaries than landscapes with different histories (mean Jaccard similarity 0.55 versus 0.35 in the 16-species community and 0.50 versus 0.28 in the three-species community). Evolution also shifted which environmental axis structured the landscape. In the 16-species community, boundaries between neighbouring streptomycin concentrations became less frequent (31.1 versus 24.8 per landscape; Holm-adjusted *P*=0.00031), whereas boundaries between neighbouring carbon levels became more frequent (16.8 versus 19.6; Holm-adjusted *P*=0.014; Fig. 2B). In the three-species community, boundaries along the streptomycin axis also became less frequent (23.4 versus 20.8; Holm-adjusted *P*=0.043), with no significant change along the carbon axis (15.1 versus 13.3; Holm-adjusted *P*=0.21). Boundaries were relocated rather than added or removed overall: the number of states did not differ significantly between histories in either community (Supplementary Table S1).

**Figure 2.**
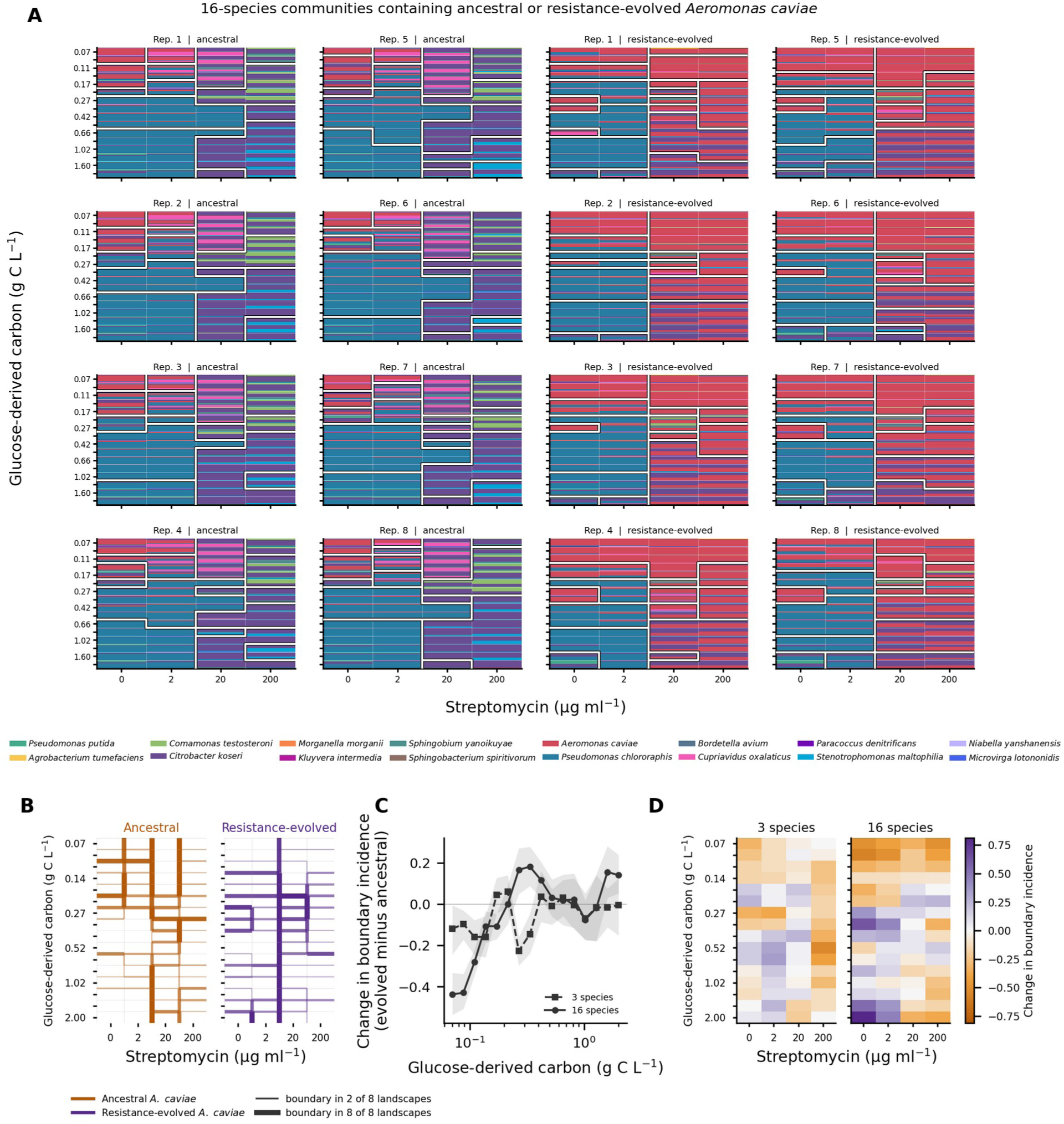
Community composition and segmented state structure across the resource-antibiotic landscape. **A** Relative abundance of species for all eight replicates of the 16-species community under ancestral and resistance-evolved *A. caviae* histories; white lines show replicate-specific segmentation boundaries. **B**, Location of community-state boundaries in 16-species landscapes containing ancestral or resistance-evolved *A. caviae*. Each edge between neighbouring environments is drawn with width and opacity proportional to the fraction of the eight replicate landscapes in which it separates two states. **C,** Evolved-minus-ancestral change in boundary incidence along the carbon gradient, averaged across streptomycin concentrations. Shading shows bootstrap 95% confidence intervals. **D**, Evolved-minus-ancestral change in mean boundary incidence across the landscape in three- and 16-species communities. Boundary incidence is the fraction of a cell’s adjacent edges that cross a state boundary.

The direction of this change depended on where in the landscape a community was (Fig. 2B–D). The history effect on boundary incidence varied across the 64 environments (exact randomization, *P*=0.00016 in both communities) and along the carbon gradient (16 species: *P*=0.00016; three species: *P*=0.0073). In the 16-species community, resistance evolution removed boundaries at low carbon (0.07–0.17 g C l^−1^; boundary incidence 0.58 versus 0.31; Holm-adjusted *P*=0.00047) and added them at intermediate carbon (0.21–0.66 g C l^−1^; 0.40 versus 0.49; Holm-adjusted *P*=0.0034). At high carbon (0.82–2.00 g C l^−1^), boundaries were gained without antibiotic and at 2 µg ml^-1^ but lost at 200 µg ml^-1^, so the net change was small (Holm-adjusted *P*=0.24). In the three-species community, boundaries were also lost at low carbon (0.42 versus 0.32; Holm-adjusted *P*=0.0065) but did not change significantly elsewhere.

These regional differences coincided with where resistance-evolved *A. caviae* reached high abundance. At low carbon, it remained abundant across streptomycin concentrations, and boundary incidence decreased relative to the ancestral history. At intermediate and high carbon, its abundance varied more strongly with streptomycin: it remained abundant at 20 and 200 µg ml^-1^ at intermediate carbon, but at high carbon it was nearly absent at 0 and 2 µg ml^-1^ and reached intermediate abundance only at 20 and 200 µg ml^-1^. Resistance evolution therefore changed where *A. caviae* was excluded, coexisted or dominated. At inhibitory concentrations, the two dominant species exchanged places. Ancestral *A. caviae* was almost absent at 20 and 200 µg ml^-1^ (mean relative abundance 0.002 and 0.004), while *C. koseri* reached 0.59 and 0.66. With resistance-evolved *A. caviae*, the focal species instead reached 0.56 and 0.67, while *C. koseri* declined to 0.30 and 0.27 (all four history contrasts Holm-adjusted *P* = 0.0006). Across individual environments, however, the change in boundary incidence was not significantly associated with how sensitive focal-species abundance was to environmental conditions (16-species community: Spearman *ρ* = 0.21, *P* = 0.25; three-species community: *ρ* = 0.22, *P* = 0.11).

### Evolution has contrasting effects on local diversity across environmental contexts

Evolutionary history did not impose a uniform change in alpha diversity (Fig. 3; Supplementary Fig. S5; Supplementary Tables S2-S3). In the three-species community, resistance evolution reduced diversity, averaged across the carbon gradient, without antibiotic (change in Shannon Index, ΔH=- 0.141) and at 2 µg ml^-1^ (ΔH=-0.128), but increased diversity at 20 and 200 µg ml^-1^ (ΔH=+0.261 and +0.450; Holm-adjusted *P*<0.001 for all four contrasts; Fig. 3A,C). The 16-species community likewise showed sign-changing responses, with ΔH=-0.168, -0.122, +0.054 and -0.091 at 0, 2, 20 and 200 µg ml^-1^, respectively (Fig. 3B,C). Thus, focal-species evolution reshaped the diversity response surface rather than shifting diversity consistently upward or downward.

**Figure 3.**
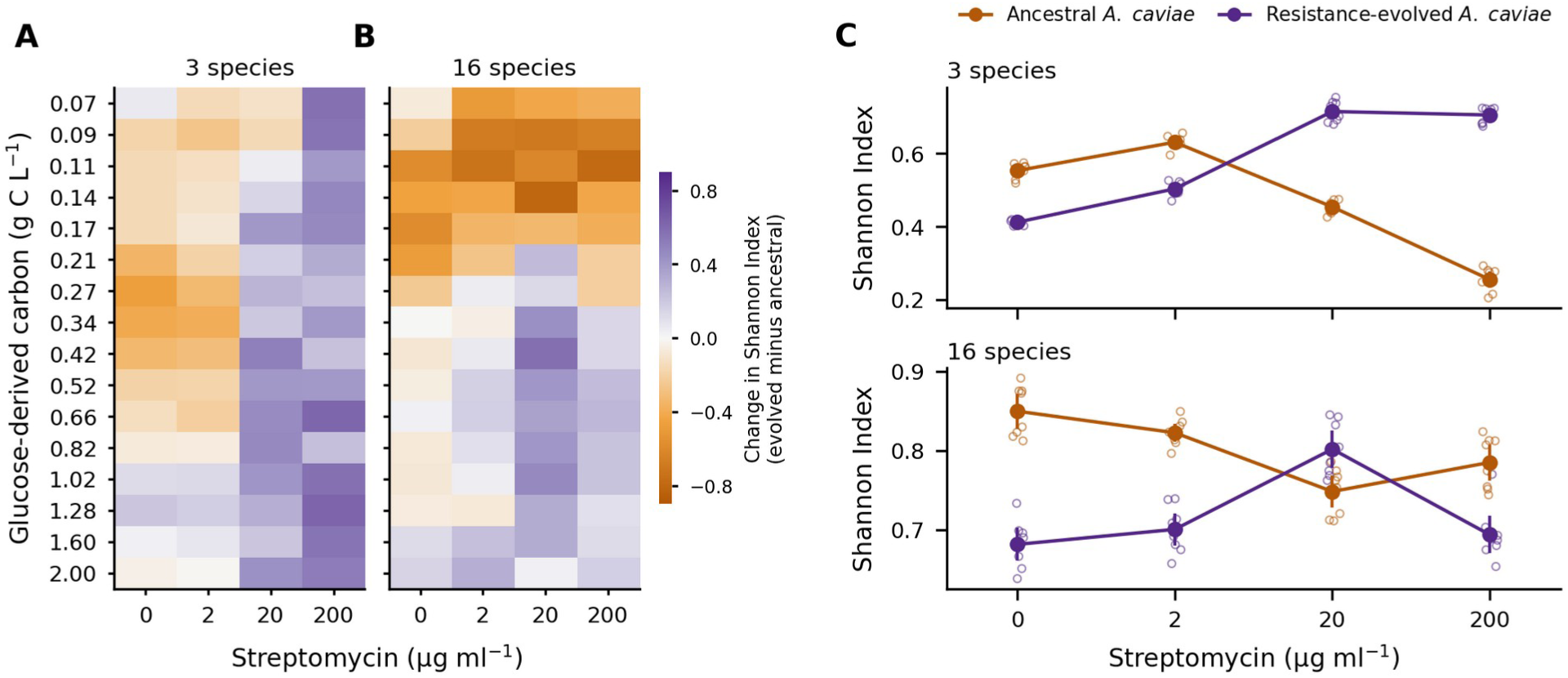
Alpha-diversity responses across the resource-antibiotic landscape. **A**,**B**, Difference in Shannon Index between resistance-evolved and ancestral focal-species histories. **C**, Shannon Index averaged across carbon at each streptomycin concentration. Open points are replicate landscapes; filled points and error bars are means and 95% confidence intervals (*N*=8 per history).

The diversity increase in the three-species community was associated with the rescue of resistance-evolved *A. caviae* in the presence of streptomycin. In the 16-species community, dominance of evolved *A. caviae* at low carbon under streptomycin was associated with reduced diversity, whereas its presence at higher carbon was associated with increased diversity. Thus, in the more complex 16-species community, resistance evolution changed whether *A. caviae* was excluded, coexisted or became dominant across environments, contributing to increases in diversity in some conditions and decreases in others.

### Evolution changes community responses far below inhibitory antibiotic concentrations

Evolutionary history altered the multivariate antibiotic response at 2 µg ml^-1^ streptomycin, below the assigned ancestral MICs of the focal strain and all members of the 16-species community (Fig. 4; Supplementary Figs. S6-S7; Supplementary Tables S4–S7). The focal species showed a strong history-dependent response at this low antibiotic concentration, and its direction depended on carbon availability (Fig. 4A,B). In the 16-species community, ancestral *A. caviae* lost 0.20–0.42 of community relative abundance from 0 to 2 µg ml^-1^ at low carbon (0.07–0.17 g C l^−1^), whereas the resistance-evolved strain changed by no more than 0.10 (history differences +0.10 to +0.49; false-discovery-rate-adjusted *q*≤0.03; Supplementary Table S8). At 0.21–0.34 g C l^−1^ the pattern reversed: the resistance-evolved strain declined by up to 0.50 while the ancestral strain increased slightly (history differences -0.16 to -0.58; *q*<0.001). Averaged across carbon, these opposite effects largely cancelled (difference +0.029; 95% CI 0.006–0.053; Holm-adjusted *P*=0.037), but the response landscapes of the two histories differed clearly (mean absolute history difference 0.19; exact *P*=0.00016; Supplementary Table S5). The three-species community showed the same protection of the resistance-evolved strain at low carbon (history differences +0.17 to +0.35; *q*≤0.02; Fig. 4A; Supplementary Table S8). This is consistent with resistance increasing its competitive fitness under sub-MIC selection^23^.

**Figure 4.**
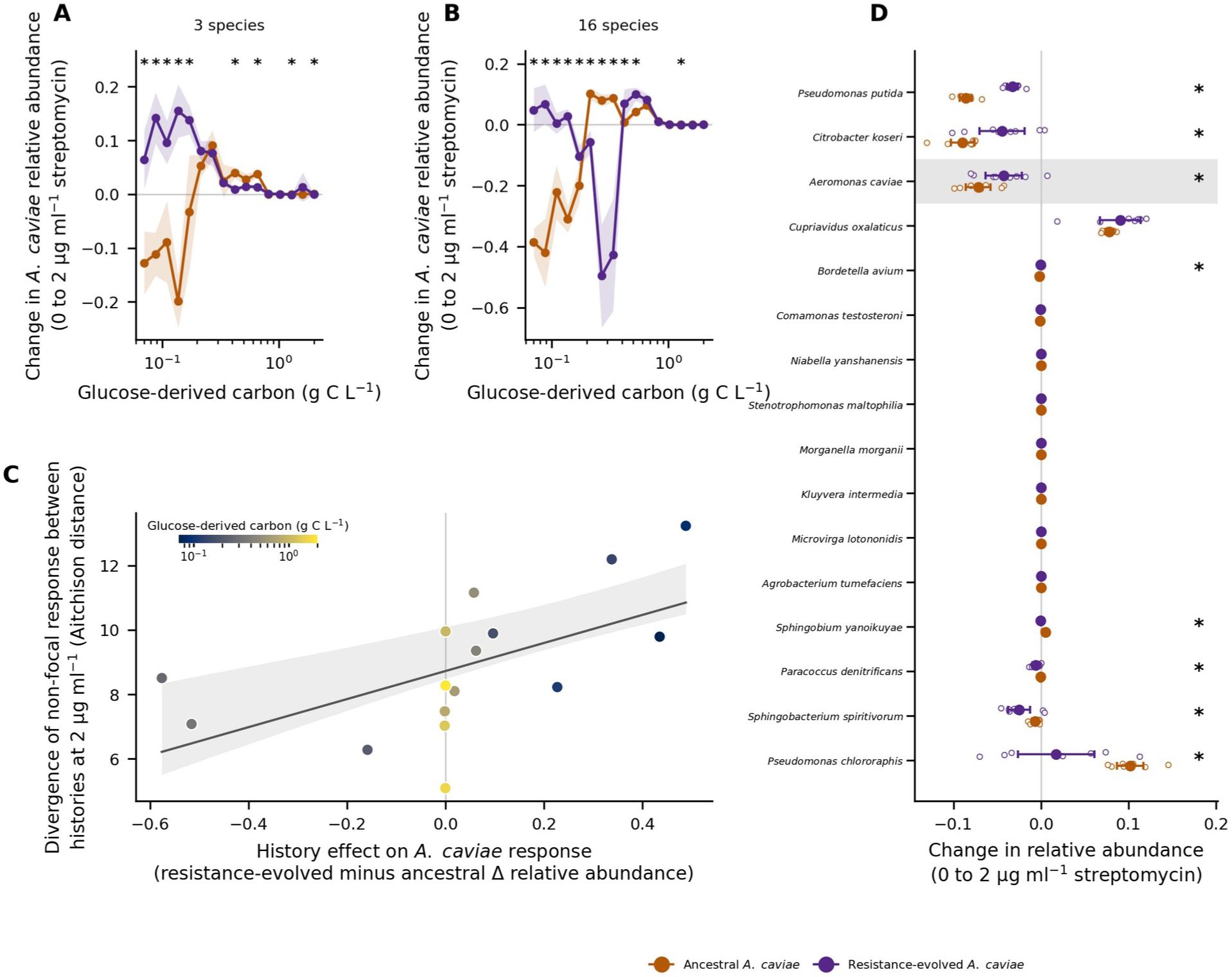
Focal-species and community responses at a low antibiotic concentration. **A,B**, Change in *A. caviae* relative abundance from 0 to 2 µg ml^-1^ streptomycin across the carbon gradient in the three-species (A) and 16-species (B) communities. Lines are means, shading 95% confidence intervals (*N*=8 per history). Asterisks mark carbon levels where the histories differ (exact allocation test, false-discovery-rate-adjusted *q* < 0.05 across 16 carbon levels). **C**, Relationship across carbon treatments between the evolutionary-history effect on the *A. caviae* response and divergence of the non-focal community response after removal of *A. caviae*. Points are carbon treatments, coloured by carbon concentration; line and shading show the fitted relationship and its bootstrapped 95% confidence intervals. **D**, Species-specific changes in relative abundance from 0 to 2 µg ml^-1^ streptomycin under each evolutionary history. *A. caviae* (shaded) is measured in the whole community, the other species within the non-focal subcomposition. Points as in A. Asterisks mark history contrasts with false-discovery-rate-adjusted *q* < 0.05 across the 15 non-focal species; *A. caviae* is marked by its carbon-averaged focal test (Holm-adjusted *P* < 0.05).

Across carbon levels, a more positive evolutionary change in the *A. caviae* response was associated with greater divergence in the response of the non-focal community (Spearman *ρ*=0.71; *P*=0.011; Fig. 4C). The non-focal response landscapes also separated between histories overall (root-mean-square Aitchison separation 9.1; exact *P*=0.00016). In addition to *A. caviae*, seven of the 15 non-focal species showed history-dependent responses after false-discovery-rate correction, three increasing and four decreasing in communities containing evolved *A. caviae* (Fig. 4D; Supplementary Table S9). Several taxa were rarely detected, and their contrasts may therefore partly reflect differences in occurrence. For example, *Paracoccus denitrificans* was detected in 10% of ancestral and 34% of resistance-evolved communities without antibiotic. Thus, resistance evolution altered the responses of other community members in different directions rather than simply displacing them as *A. caviae* increased. This is consistent with indirect competitive effects, whereby suppression of some species by *A. caviae* can release others from competition^19,24^. Because relative abundances sum to one, losses by some species at the low antibiotic concentration were balanced by gains in others. Under the ancestral history, *Pseudomonas chlororaphis* and *Cupriavidus oxalaticus* gained most of the non-focal abundance lost by *Pseudomonas putida* and *C. koseri* (Fig. 4D). Resistance evolution in *A. caviae* weakened the exchange between the two *Pseudomonas* species. With resistance-evolved instead of ancestral *A. caviae*, the decrease in *P. putida* relative abundance from 0 to 2 µg ml^−1^ streptomycin was smaller (0.032 instead of 0.086), as was the increase in *P. chlororaphis* (0.017 instead of 0.102). In contrast, the gain of *C. oxalaticus*, the only member of the Burkholderiales among the abundant species, did not differ between histories (Supplementary Table S9). The species that gained were not the most resistant. *C. koseri*, which had the highest MIC of the five abundant species, lost ground, whereas *P. chlororaphis*, with one of the lowest MICs, gained. Thus, evolution of the focal species mainly changed the competitive outcome between two closely related species that are likely to occupy similar niches. The same two pairs (*P. chlororaphis* with *P. putida*, and *A. caviae* with *C. koseri*) also replaced each other in the long-term experiment from which the evolved *A. caviae* clone originated^19^.

### Resistance evolution changes which traits explain species success

We next asked whether species success across the landscape could be explained by two kinds of traits measured in monocultures: intrinsic streptomycin resistance and growth traits. Antibiotic exposure enriched intrinsically resistant taxa in the 16-species community after excluding resistant *A. caviae* (Fig. 5A; Supplementary Figs. S8-S9; Supplementary Tables S10-S11). In communities containing ancestral *A. caviae*, high-MIC taxa carried 26% of non-focal abundance without antibiotic, increasing to 69% at 20 µg ml^-1^ and 85% at 200 µg ml^-1^. The corresponding values with resistance-evolved *A. caviae* were 19%, 78% and 80%. Relative to the antibiotic-free condition, enrichment at 20 and 200 µg ml^-1^ occurred in both histories (paired Holm-adjusted *P*=0.023). Differences between histories were small, at most 9 percentage points, and their direction varied with dose (Supplementary Table S10). Resistance evolution thus changed resistance-based sorting among the other species only slightly. Instead, it made *A. caviae* part of the resistant group, where it competed directly with the intrinsically resistant taxa. *C. koseri*, the most abundant of these, declined from 0.66 to 0.27 of community relative abundance at 200 µg ml^-1^.

**Figure 5.**
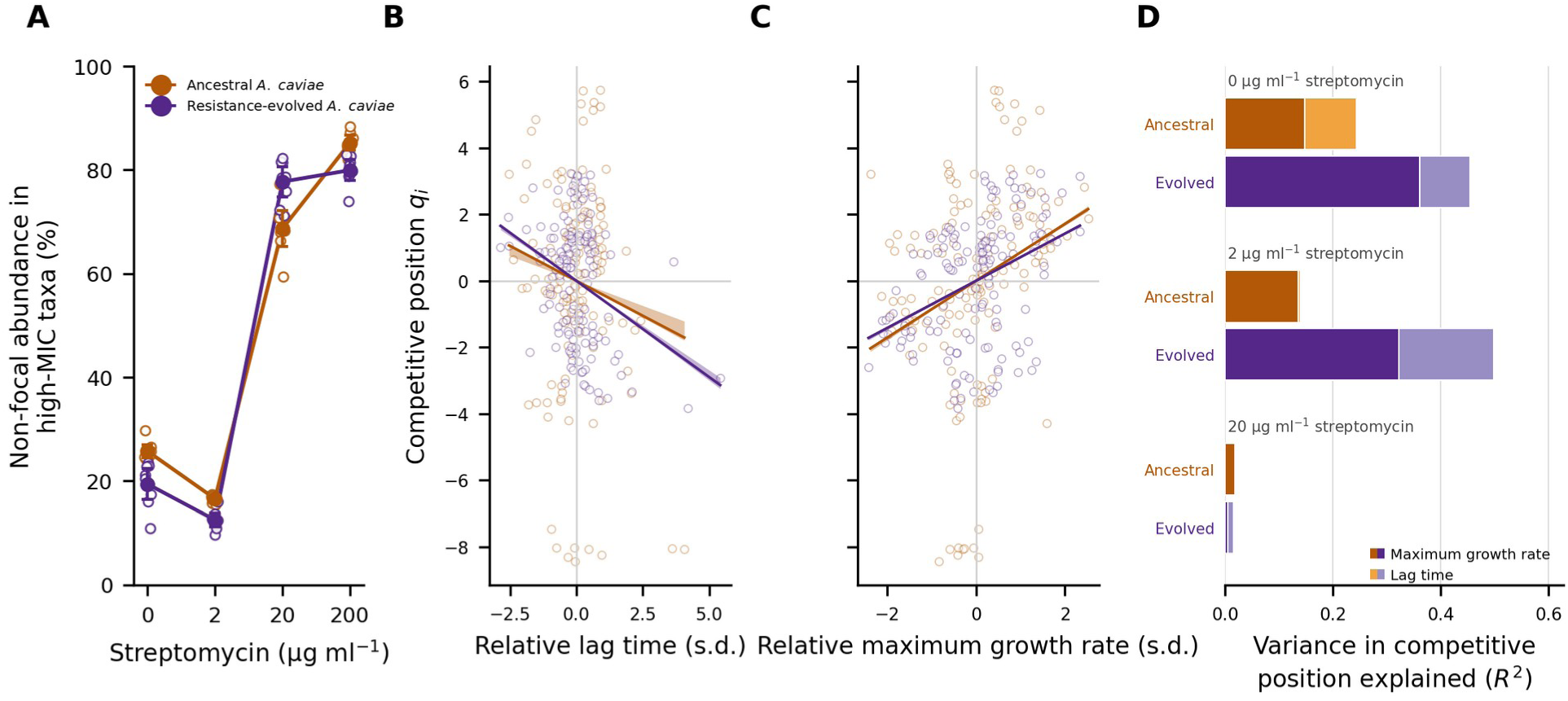
Traits associated with species success across the antibiotic gradient. **A**, Fraction of non-focal abundance carried by high-MIC taxa in 16-species communities. Open points are replicates; filled points and error bars show means and 95% confidence intervals (*N*=8 replicates per history). **B,C**, Relationship between relative monoculture lag time (B) or maximum growth rate (C) and endpoint competitive position *q_i_* in three-species communities. *q_i_* is the centred-log-ratio coordinate of a species’ endpoint abundance, with larger values indicating greater abundance relative to the other community members. Points are species × carbon × streptomycin condition means, based on *N*=6 monoculture growth replicates and *N*=8 community replicates per condition; lines show the median slope across the *N*=8 replicate landscapes per history from linear models including both growth traits, with shading showing its bootstrap 95% confidence interval. Points near −8 are ancestral *A. caviae* at 1.02–1.60 g C l^-1^, where it was excluded from the community. **D**, Dose-resolved Shapley/LMG partition of the *R*^2^ from the same two-trait models. Stacked bars give the variance explained by each trait; total bar length gives the joint *R*^2^. Values are means across *N*=8 replicate-specific fits per history. Panels B-D exclude 200 µg ml^-1^ streptomycin because kinetic traits were unreliable under negligible growth.

We then investigated whether monoculture growth traits explained the realized competitive position of the three dominant species across the environmental landscape. For each environment, maximum growth rate and lag time were expressed relative to the mean of the three community members and related to each species’ endpoint centred-log-ratio coordinate, *q_i_*, where higher *q_i_* indicates greater abundance relative to the other community members (Fig. 5B-D; Supplementary Figs. S10-S12; Supplementary Tables S12-S14). Analyses of growth kinetics were restricted to 0, 2 and 20 µg ml^-1^ streptomycin because negligible growth of ancestral *A. caviae* and *P. chlororaphis* at 200 µg ml^-1^ made species comparisons impossible, with only *C. koseri* remaining in some environments.

Using replicate-stratified two-predictor linear models, maximum growth rate was positively associated with competitive position under both focal-species histories (mean partial slope 0.865 under the ancestral history and 0.717 under the resistance-evolved history; medians 0.856 and 0.710; Holm-adjusted *P*=0.0156 for both), although the association was weaker after focal-species evolution (evolved-minus-ancestral slope difference -0.148; Holm-adjusted *P*=0.00031). Lag time showed a similar history dependence, with a negative association under the ancestral history (slope -0.399; median -0.420; Holm-adjusted *P*=0.0156) that was stronger under the resistance-evolved history (-0.547; median -0.579; Holm-adjusted *P*=0.0156; history difference -0.148; Holm-adjusted *P*=0.0064). Without antibiotic, maximum growth rate and lag time jointly explained 24-45% of variation in competitive position, whereas at 2 µg ml^-1^ maximum growth rate accounted for 98% (ancestral history) and 66% (resistance-evolved history) of the variation explained by these two traits. At 20 µg ml^-1^, the two traits together explained little variation in competitive position.

Thus, without antibiotic, competitive ranking reflected both how early species entered growth and how rapidly they subsequently grew. At low streptomycin, under the ancestral history, the contribution of lag largely disappeared while maximum growth rate remained associated with competitive position. This is consistent with streptomycin changing the selective meaning of lag: delayed growth reduces the competitive advantage of early resource capture, but can also delay entry into the metabolically active state in which aminoglycoside effects are strongest. Under the resistance-evolved history, lag time still accounted for about one third of the explained variation at this concentration (Supplementary Table S14). This reflects the altered competitive ranking in communities containing the evolved strain. At higher antibiotic concentrations, susceptible species declined or disappeared, leaving fewer effective competitors and consequently less scope for either growth trait to explain competitive ranking. Species success was therefore partly explained by traits measured in monocultures. However, the relevant traits changed along the antibiotic gradient, from growth traits at zero and low streptomycin to intrinsic resistance at inhibitory concentrations. By changing which species dominated, resistance evolution in *A. caviae* changed which traits were associated with success.

## Discussion

Our results show that evolution in a single species can reorganize how entire communities respond to environmental conditions. Resistance evolution changed where community-state boundaries were located across the resource-antibiotic landscape, removing boundaries at low carbon, where the focal species dominated at every antibiotic concentration, and adding them at intermediate carbon, where its abundance depended on streptomycin. It also increased diversity in some environments while decreasing it in others. Previous work has shown that evolution can alter community structure^1,4,19^ and that the effects of antibiotic resistance depend on context^14–17^. Our results connect these two lines of work: the community consequences of the same evolutionary change depended on the environment in which they were measured.

These community changes can be traced to how resistance evolution changed *A. caviae*. At most carbon levels, the evolved strain started growing later but then grew slightly faster, and only the evolved strain could grow at the highest streptomycin concentration. These changes altered competition between two pairs of closely related species. At inhibitory streptomycin concentrations, evolved *A. caviae* replaced *C. koseri* as the most abundant species. Without antibiotic, *P. putida* and *P. chlororaphis* coexisted in communities with ancestral *A. caviae*, and low streptomycin shifted abundance from *P. putida* to *P. chlororaphis*. Evolved *A. caviae* was more abundant at most low and intermediate carbon levels, and in its presence *P. putida* was nearly lost even without antibiotic, so this shift largely did not occur (Fig. 4D). These changes in which species won explain much of the change in community-state boundaries and diversity.

Resource availability and antibiotic stress played different roles in this reorganization. Resource availability determined where community-state boundaries were lost or gained, while antibiotic concentration determined the direction of change at high carbon. Carbon availability even reversed how resistance evolution affected the focal species at low streptomycin: the evolved strain was protected from the loss in abundance suffered by the ancestor when carbon was scarce, but declined more than the ancestor at slightly higher carbon. The more resistance evolution improved the response of *A. caviae*, the more the response of the rest of the community diverged between histories (Fig. 4C). Environmental stress is known to influence community assembly^5^, and selection for antibiotic resistance depends on environmental conditions^25^. Our results show that resource availability also determines how resistance evolution, once it has occurred, affects the rest of the community. The resource gradient in our experiments was represented by glucose. It has recently been shown that dietary sugar also modifies the community effects of antibiotics in a clinical context^26^.

Both pairs of species whose balance changed are closely related and likely to overlap in resource use^19^. The indirect effects of evolution in this community may therefore act mainly through competition between species with similar niches, where some taxa were suppressed while others may have experienced ecological release^19,24^. These effects also occurred at sub-MIC streptomycin concentrations, showing that the ecological consequences of resistance evolution do not require direct growth inhibition of all community members^27,28^. Because community composition reflects the outcome of competition rather than its course, these patterns identify which competitive relationships changed, while the underlying interaction mechanisms remain an open question.

Species success was partly explained by traits measured in monocultures, but which traits mattered depended on the environment and on evolutionary history. When all dominant species could grow, both lag time and maximum growth rate were associated with competitive sorting. At low streptomycin, maximum growth rate became the more important trait, whereas at higher concentrations, susceptible competitors were increasingly excluded and resistance became more important for competitive position. This held in both histories, but resistance evolution moved *A. caviae* into the resistant group (Fig. 5A), which also changed which growth traits explained competitive position at low streptomycin. Lag time and growth rate are known to influence microbial competition^29^, environmental conditions modify their importance^21^, and competitive outcomes also depend on resource competition and antibiotic conditions^14^. The trait relationships we observed are correlative, and part of the variation in competitive position likely reflects interactions that monoculture traits do not capture. The effects of the evolved strain also reflect its whole evolved phenotype rather than resistance alone, as is the case when evolved lineages spread in natural communities.

Evolution in a single species can change not only the composition of a community but also how it responds to its environment. Because such effects can reverse along environmental gradients, studies in a single environment may misjudge both their size and their direction. These effects are nevertheless not arbitrary. Traits measured in monoculture, together with competition between closely related species, can partly explain how evolution reshapes community responses. Accounting for environmental context and species traits together may improve our understanding of community responses and resilience under anthropogenic disturbance^10,22,24,30^, including the consequences of antibiotic resistance evolution.

## Methods

### Bacterial species and experimental communities

We used a previously established 16-species synthetic bacterial community described by Bischofberger et al. (2026)^21^. All ancestral species were obtained from the HAMBI culture collection (University of Helsinki, Finland). The community consisted of *Pseudomonas putida* (HAMBI 6), *Agrobacterium tumefaciens* (HAMBI 105), *Comamonas testosteroni* (HAMBI 403), *Citrobacter koseri* (HAMBI 1287), *Morganella morganii* (HAMBI 1292), *Kluyvera intermedia* (HAMBI 1299), *Sphingobium yanoikuyae* (HAMBI 1842), *Sphingobacterium spiritivorum* (HAMBI 1896), *Aeromonas caviae* (HAMBI 1972), *Pseudomonas chlororaphis* (HAMBI 1977), *Bordetella avium* (HAMBI 2160), *Cupriavidus oxalaticus* (HAMBI 2164), *Paracoccus denitrificans* (HAMBI 2443), *Stenotrophomonas maltophilia* (HAMBI 2659), *Niabella yanshanensis* (HAMBI 3031) and *Microvirga lotononidis* (HAMBI 3237).

We created four community types that differed in species richness and in the evolutionary history of HAMBI 1972. The three-species ancestral community consisted of ancestral HAMBI 1972, HAMBI 1977 and HAMBI 1287. The corresponding evolved community consisted of streptomycin-evolved HAMBI 1972, ancestral HAMBI 1977 and HAMBI 1287. The 16-species ancestral community consisted of ancestral HAMBI 1972 together with the other 15 ancestral species. The evolved 16-species community contained the same evolved HAMBI 1972 together with the same 15 ancestral species. Thus, the ancestral and evolved versions of each community differed only in the evolutionary history of HAMBI 1972.

### Origin and phenotypic characterization of evolved HAMBI 1972

The streptomycin-evolved HAMBI 1972 clone used in this study (EVO1972 3035 BS) originated from the long-term microbial community evolution experiment described by Kivikoski et al. (2026)^19^. The clone was isolated at experimental time point 151, that is after 151 weekly serial transfers (approximately 2.9 years of evolution), from a community grown in an environment containing 20 µg ml^−1^ streptomycin. In this experiment, streptomycin resistance in *A. caviae* arose through a selective sweep of the *rpsL* point mutation Lys88Arg, which was confirmed by whole-genome sequencing of resistant clones isolated at the same time point^19^.

To isolate evolved strains, samples from the evolved communities were plated on CHROMagar, and individual colonies were picked and cryopreserved in 96-well glycerol stocks. Candidate HAMBI 1972 clones were phenotypically screened across a streptomycin concentration gradient. Growth was assayed at 0, 1.25, 2.5, 5, 10, 20, 40, 80 and 160 µg ml^−1^ streptomycin. EVO1972 3035 BS grew at the highest tested streptomycin concentration of 160 µg ml^−1^. Without streptomycin, the evolved and ancestral clones grew similarly in monoculture. Differences in growth traits depended on carbon level (Supplementary Figs. S10 and S12). For example, yield was within 10% of the ancestral value at every carbon level. Lag time, however, was longer at 13 of 16 carbon levels, by 0.6 h on average, suggesting a growth cost of streptomycin resistance. Such changes are expected outcomes of resistance evolution and may also reflect adaptation to the growth medium.

### Growth medium and carbon gradient

Experimental communities were grown in M9 minimal medium with glucose as the sole added carbon source. M9 minimal medium was prepared by dissolving 11.28 g of M9 minimal medium salts (MP Biomedicals, UK) in 1 l of Milli-Q purified water. The medium was autoclaved and supplemented with filter-sterilized MgSO_4_ and CaCl_2_ stock solutions to final concentrations of 2 mM and 0.1 mM. Glucose was added from a filter-sterilized 500 g l^−1^ stock solution.

Resource availability was manipulated using a 16-level glucose-derived carbon gradient. The gradient was generated by serial 0.8-fold dilution. The resulting carbon concentrations were 0.0704, 0.0880, 0.1100, 0.1374, 0.1718, 0.2147, 0.2684, 0.3355, 0.4194, 0.5243, 0.6554, 0.8192, 1.024, 1.28, 1.60 and 2.00 g C l^−1^.

### Streptomycin treatments

Antibiotic stress was manipulated using four final streptomycin concentrations: 0, 2, 20 and 200 µg ml^−1^. The four streptomycin levels were fully crossed with the 16 carbon concentrations, resulting in 64 distinct combinations of carbon availability and antibiotic concentration.

### Preculturing and community assembly

Individual bacterial species were grown from frozen stocks in 6 ml of proteose peptone yeast extract medium (PPY; 20 g proteose peptone and 2.5 g yeast extract per litre of deionized water) for 48 h at 30 °C with constant shaking at 70 rpm. A 5-ml aliquot of each culture was centrifuged for 5 min at 3,000 × g at room temperature. The supernatant was removed, and the cells were washed with 3 ml M9 medium and centrifuged for 3 min at 3,000 × g. The washed pellets were resuspended in M9 medium.

Cell density was estimated as optical density at 600 nm (OD_600_) using an Absorbance 96 plate reader (Byonoy), with M9 medium as a blank, and cultures were standardized to OD_600_ = 1.4. Communities were assembled by combining equal volumes of the standardized cultures.

### Experimental design

We combined two levels of community species richness (3 or 16 species), two evolutionary histories of HAMBI 1972 (ancestral or evolved), 16 carbon concentrations and four streptomycin concentrations. Each combination of community type, carbon concentration and streptomycin concentration had eight independent replicate populations, resulting in 2,048 experimental populations in total.

For each community type, the 64 combinations of carbon and streptomycin concentrations were arranged on six 96-well deep-well plates. The eight replicate populations for each treatment were located in rows A–H and were transferred independently throughout the experiment. The same plate layout was used for each community type. Uninoculated wells were included as negative controls to monitor contamination.

### Serial-passage community experiment

Each assembled community was inoculated into 96-well deep-well plates by adding 10 µl of community culture (approximately 1% inoculum) to 1 ml of M9 minimal medium containing the assigned carbon and streptomycin concentrations. Plates were sealed with breathable sealing films and incubated for 48 h at 30 °C with continuous shaking at 800 rpm. After each 48-h growth period, 10 µl of culture (approximately 1% of the culture volume) from each population was transferred into the corresponding well of a fresh plate containing the same carbon and streptomycin treatment. Each replicate population was transferred independently throughout the experiment. Eight serial transfers were performed at 48 h intervals. The experiment lasted 16 days.

### Growth measurements and sample storage

At the end of each 48-h growth period, 100 µl of culture from each experimental population was transferred to a separate measurement plate, and growth was quantified as OD_600_. After each transfer, plates from the completed growth cycle were sealed with adhesive aluminum foil and stored at −20 °C for 16S rRNA gene amplicon sequencing of community composition.

### Monoculture growth assay

Ancestral HAMBI 1972, HAMBI 1287 and HAMBI 1977 and the streptomycin-evolved HAMBI 1972 (EVO1972 3035 BS) were grown separately to characterize monoculture growth across the carbon and streptomycin gradients. Cultures were grown from frozen stocks in 6 ml PPY medium for 48 h at 30 °C with constant shaking at 70 rpm. A 500-µl aliquot of each culture was centrifuged for 5 min at 3,000 × *g* at room temperature. The supernatant was removed, and the cell pellet was washed with 500 µl M9 medium and centrifuged for 3 min at 3,000 × *g*. The supernatant was removed, and the pellet was resuspended in 500 µl M9 medium. Each cell suspension was diluted 1:10,000 in M9 medium.

Ten microlitres of each diluted cell suspension was transferred into wells of 96-well plates containing 140 µl of glucose- and streptomycin-supplemented M9 minimal medium corresponding to the carbon and streptomycin treatments described above. Each combination of monoculture, carbon concentration and streptomycin concentration had six independent replicates.

Plates were incubated in LogPhase 600 Microbiology Readers (Agilent BioTek, Santa Clara, CA, USA) at 30 °C for 48 h with shaking at 800 rpm. Optical density at 600 nm (OD_600_) was measured every 10 min. At the end of each 48-h growth period, 10 µl of culture from each monoculture was transferred into the corresponding well of the next transfer plate containing the same carbon and streptomycin treatment. Three 48-h growth cycles were performed.

### 16S rRNA gene amplicon sequencing

Community composition was quantified by sequencing 16S rRNA gene amplicons, using the protocol established for this community^19,24^. DNA was extracted from frozen samples collected at the final timepoint (day 16) using the DNeasy 96 Blood & Tissue Kit (Qiagen). NanoDrop (Thermo Scientific) and the Quant-iT 1X dsDNA HS kit (Invitrogen) were used to measure sample quality and quantity.

The TaggiMatrix multiplexing protocol was used for amplicon library preparation^31^. First, the V3 region of the 16S rRNA gene was amplified using primers containing iTru fusion adapters (Fwd 5′-CCTACGGGAGGCAGCAG-3′, Rev 5′-ATTACCGCGGCTGCTGG-3′) and a sample-specific combinatorial index. The NGS Normalization 96-Well Kit (Norgen Biotek) was used for purification. Up to 96 samples were pooled and subjected to limited-cycle PCR using standard Illumina TruSeq primers and indices. The reaction products were purified using Sera-Mag particles (Cytiva) and then pooled equimolarly. The Bioanalyzer High Sensitivity DNA Kit (Agilent) was used for library quality control. Paired-end sequencing (2 × 150 bp reads) was performed at the Finnish Functional Genomics Centre, Turku, Finland, using the Illumina MiSeq 2000 V2 platform.

### Sequence analysis

Reads were processed with the bioinformatic pipeline established for this community^19,24^. Demultiplexing proceeded in two stages, first on the i5/i7 index pairs in Illumina BaseSpace and then on the custom TaggiMatrix indices. Primer sequences were trimmed with cutadapt^32^ version 4.4.

Forward and reverse reads were joined into single amplicons with NGmerge^33^ version 0.3.0. Quality filtering used vsearch^34^ version 2.22.1. Reads were discarded if they contained ambiguous bases, carried more than one expected error, or fell outside 115–165 bp. Surviving reads were assigned to strains with Rbec^35^ version 1.8.0 with a reference database consisting of unique V3 amplicon sequences from the 16 HAMBI bacterial species. To correct for differences in 16S rRNA gene copy number between strains, counts were scaled by the number of full-length 16S rRNA genes per genome matching the V3 primer pair^36,37^. Computer code for the workflow is available at gitlab.utu.fi/slhogl/hambiAmplicon.

### Downstream analysis

Analyses were performed in Python 3.13.5 in Jupyter notebooks using NumPy, pandas, SciPy, statsmodels and Matplotlib. Full definitions and inferential procedures are provided in the Supplementary Methods. Endpoint read counts were converted to normalized species relative abundances. Each carbon × streptomycin × focal-species-history condition contained *N*=8 independently cultured replicate communities. For analyses requiring a complete 16-carbon × 4-streptomycin response surface, eight analytical landscapes were constructed by fixed replicate-index assignment: replicate 1 from every environmental condition was assigned to landscape 1, through to replicate 8 assigned to landscape 8. This arbitrary but fixed grouping ensured that each independent community entered each landscape analysis once and avoided pseudoreplication from enumerating multiple cross-condition replicate combinations. The same constructed landscapes were retained throughout all analyses requiring a complete response surface. Shannon Index was calculated from non-zero proportions. Community-state landscapes were segmented independently for each constructed landscape using a custom greedy spatial algorithm that merged adjacent cells when this reduced the sum of within-state generalized Jensen-Shannon compositional loss and a boundary penalty (for details, see Supplementary Methods). Merges were constrained to a maximum within-state Bray-Curtis diameter of 0.40; the baseline boundary penalty was 0.15 and sensitivity analyses varied both parameters^38,39^.

For each environment, we quantified the sensitivity of resistance-evolved *A. caviae* abundance to environmental conditions as the magnitude of its local gradient across the carbon × streptomycin grid. We related this measure to the evolved-minus-ancestral change in community-state boundary incidence using Spearman correlation, separately for three- and 16-species communities. Significance was assessed by exact randomization of evolutionary-history labels across the 16 constructed landscapes (6,435 distinct allocations), recalculating both measures for each allocation. Full procedures are described in the Supplementary Methods.

Changes in landscape topology were analysed on the state boundaries of the segmented landscapes. Each constructed landscape was represented by its 108 adjacent-cell edges (60 along the carbon axis and 48 along the streptomycin axis), coded 1 where an edge separated two states. Global differences in boundary configuration between focal-species histories were tested by exact two-group PERMANOVA on Euclidean distances between these vectors. As a sensitivity analysis, whole partitions were compared using one minus the adjusted Rand index. Boundary orientation was summarized as the number of boundary edges per landscape along each axis and compared between histories by exact allocation tests, with Holm correction across the two axes. The boundary incidence of an environment was the fraction of its adjacent edges that crossed a state boundary. Spatial heterogeneity of the history effect was tested with the sum of squared deviations of the evolved-minus-ancestral change in mean boundary incidence from its landscape mean, over the 64 environments or over the 16 carbon levels after averaging across streptomycin, using exact allocation randomization. Regional contrasts compared low (0.07–0.17 g C l^−1^), intermediate (0.21–0.66 g C l^−1^) and high (0.82–2.00 g C l^−1^) carbon, with Holm correction across the three regions. At 2 µg ml^-1^ streptomycin, the focal response was also compared between histories at each carbon level by exact allocation tests with Benjamini-Hochberg correction across the 16 levels, and summarized as the mean absolute history difference across carbon levels, tested by the same allocation randomization.

Compositional antibiotic responses were analysed after adding a 10^-6^ pseudocount and applying a centred-log-ratio (CLR) transformation, using the matched no-antibiotic condition at the same carbon concentration as baseline. Species-level responses at the low dose are reported as changes in relative abundance. *A. caviae* is absent from part of the ancestral landscape but from none of the resistance-evolved landscape, so log-ratio coordinates of its abundance would depend on the constant used to replace zeros; log-ratio versions of these contrasts, and their sensitivity to that constant, are given in Supplementary Table S7 and the Supplementary Methods. *A. caviae* is measured in the whole community and the other species within the non-focal subcomposition, so that their changes are not forced by the focal species’ change. Whole-community multivariate analyses retain all 16 species. For a composition, *p_i_* denotes the relative abundance of species *i*, while *p_j_* denotes the relative abundance of species *j* and *S* the number of species included in the composition; the CLR coordinate of species *i*, denoted *q_i_* in the three-species analyses, was defined as

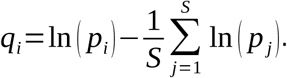

Euclidean distances between CLR vectors were interpreted as Aitchison distances^40^. Resistance filtering was quantified as the non-focal abundance carried by taxa with ancestral streptomycin MIC ≥160 µg ml^-1^. Monoculture growth curves were summarized as maximum growth rate, lag time, yield and growth-curve AUC from six replicates per strain × carbon × streptomycin condition. For the three-species communities, q*_i_* was the CLR coordinate defined above and was used as endpoint competitive position; higher q*_i_* indicates greater abundance relative to the geometric mean abundance of the three species. Kinetic-trait analyses were restricted to 0, 2 and 20 µg ml^-1^ streptomycin because ancestral *A. caviae* and *P. chlororaphis* showed negligible growth at 200 µg ml^-1^. Let r*_i_* and ℓ*_i_* denote maximum growth rate and lag time of species i, and r^\^ and ℓ\ their three-species means within the same environment. Relative traits were centered on these means and standardized. For each of the eight constructed community landscapes per history, the following two-predictor ordinary least-squares model was fitted:

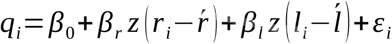

The coefficients *β_r_* and *β_ℓ_* therefore estimate the partial association of maximum growth rate and lag time with competitive position while controlling for the other trait. The eight coefficients from the fixed constructed landscapes per focal-species history were the inferential observations. Figures show the median of the eight coefficients. Dose-specific models used the same formulation, and joint model *R*^2^ was partitioned between the two traits using the two-predictor Shapley/LMG decomposition.

Individual community cultures were the biological replicates at each environmental condition. For analyses that summarized or modelled complete response surfaces, the eight fixed constructed landscapes per history defined above were used as the analytical units, so that every underlying community contributed once to each landscape-level statistic. For univariate ancestral-versus-evolved contrasts, the test statistic was the difference in constructed-landscape means and two-sided exact allocation randomization evaluated all 12,870 assignments of eight ancestral and eight evolved labels. Within-history contrasts based on the same fixed construction, and within-history tests of growth-trait slopes, used exact sign-flip randomization of the eight constructed-landscape contrasts or slopes; history differences in growth-trait slopes used exact allocation randomization of the 16 slope estimates.

Global landscape effects were tested by exact two-group PERMANOVA, representing each constructed landscape by its 64 Shannon Index values or relevant CLR response coordinates and using pseudo-*F* as the test statistic^41^. The low-dose focal/non-focal association used Spearman ρ with evolutionary-history labels randomized across the 16 constructed landscapes. Exact *P* values were the proportion of permitted randomizations at least as extreme as the observed statistic, and confidence intervals were obtained from 10,000 bootstrap resamples of the eight constructed landscapes within history. Condition-level summaries and figures retain N=8 independent biological community replicates per treatment. Holm correction was used for prespecified small test families and Benjamini-Hochberg correction for species-wide scans^42,43^.

## Author contributions

Inga-Katariina Aapalampi (Investigation, Methodology, Writing – original draft, Writing – review & editing, Funding acquisition), Anna M Bischofberger (Investigation, Resources, Writing – review & editing), Dovydas Kičiatovas (Formal analysis, Visualization, Writing – review & editing), Mikko Kivikoski (Formal analysis, Visualization, Writing – review & editing); Ville Mustonen (Conceptualization, Visualization, Investigation, Supervision, Writing – review & editing), Teppo Hiltunen (Conceptualization, Visualization, Investigation, Supervision, Funding acquisition, Writing – review & editing); Johannes Cairns (Conceptualization, Formal analysis, Visualization, Investigation, Methodology, Supervision, Funding acquisition, Writing – original draft, Writing – review & editing).

## Acknowledgements

We thank Fanny Koskela, Linda Nevala, Sanna Pausio, Hilla Pellikka, Julia Saloranta and Irma Warinowski for assistance with the laboratory work.

## Data availability

The 16S rRNA amplicon sequencing data from the serial-passage experiment are available in the European Nucleotide Archive (ENA) under accession number PRJEB129114. Downstream community composition and monoculture growth data are available at github.com/johannescairns/evolution-landscape and archived on Zenodo (https://doi.org/10.5281/zenodo.24369115).

## Code availability

Computer code for the 16S rRNA amplicon sequencing workflow is available at gitlab.utu.fi/slhogl/hambiAmplicon. The code used for the downstream analyses and figures is available at github.com/johannescairns/evolution-landscape and archived on Zenodo (https://doi.org/10.5281/zenodo.24369115).

## Use of AI

Generative AI tools were used to assist with editing language and code in the manuscript. All AI-assisted text and code was reviewed and approved by the authors.

