## Supplementary Information for "Resistance evolution in a single species reorganizes community responses across a resource-antibiotic landscape"

**This file includes**

- Supplementary Methods
- Supplementary Figures S1-S12
- Supplementary Tables S1-S14

### Supplementary Methods

#### Data structure and compositional quantities

Each carbon × streptomycin × focal-species-history treatment contained $N$=8 independently cultured biological replicate communities. To analyse complete 16-carbon × 4-streptomycin response surfaces without generating and treating multiple cross-condition replicate combinations as additional replicates, eight analytical landscapes were constructed by fixed replicate-index assignment: replicate 1 from every condition was assigned to landscape 1, through to replicate 8 assigned to landscape 8. This grouping was therefore an arbitrary but fixed analytical construction, not a biological block. The same eight constructed landscapes were used consistently for segmentation, landscape-level summaries, multivariate landscape tests, low-dose cross-carbon coupling and replicate-stratified growth-trait analyses. Each underlying biological community thus entered a given analysis only once.
For a composition containing $S$ species with relative abundances $p_i$, Shannon Index was calculated from non-zero proportions as:

$$H = -\sum_{i=1}^{S} p_i \ln(p_i)$$

Compositional antibiotic responses used a $10^{-6}$ pseudocount, after which proportions were reclosed and centred-log-ratio (CLR) transformed. Here $p_i$ is the relative abundance of species $i$, $p_j$ is the relative abundance of species $j$, and $S$ is the number of species retained in the composition:

$$clr_i(p) = \ln(p_i) - \frac{1}{S}\sum_{j=1}^{S} \ln(p_j)$$

ΔCLR denotes the CLR coordinate at a given antibiotic concentration minus the matched no-antibiotic CLR coordinate at the same carbon concentration. Species-level responses at the low dose are reported as changes in relative abundance rather than as log-ratio coordinates. Seven of the 16 taxa are detected in only 0.3-17.6% of samples, and *A. caviae* itself is absent from 12.5% of ancestral no-antibiotic conditions and 8.6% of ancestral conditions at 2 µg $ml^{-1}$, and from none under the resistance-evolved history. Any log-ratio coordinate of a taxon that is structurally absent in one arm and not the other is therefore governed by the constant used to replace zeros: the evolved-minus-ancestral CLR contrast for the focal species moves from -0.94 through -0.74 to -0.55 as the pseudocount is raised from $10^{-6}$ through $10^{-4}$ to $10^{-3}$, and the number of taxa called significant ranges from three to eleven. Restricting the basis to taxa detected in more than half of samples does not remove the dependence, because several of those taxa are themselves undetected in a third to a half of samples. Relative abundance needs no zero replacement. *A. caviae* is measured in the whole community; the other species are measured within the non-focal subcomposition, with *A. caviae* removed and the remainder reclosed, so that their changes are not an arithmetic consequence of its change. *A. caviae* cannot be placed on that basis because the basis is defined by its removal. Contrasts for the 15 non-focal species were corrected together (Benjamini-Hochberg); the focal contrast is the primary focal-response test (Supplementary Table S5). The whole-community version for all 16 species gives 6 of 16 significant contrasts, three increasing and three decreasing; both versions are tabulated in the analysis archive (Supplementary Table S9 and results/statistics/lowdose_species_contrasts_whole_community.csv in the code repository), and the log-ratio version on the prevalence-filtered basis is given in Supplementary

Table S7. Whole-community multivariate analyses retain all 16 species. Aitchison distance between two compositions x and y was calculated as Euclidean distance between their CLR vectors:

$$d_A(x, y) = \| clr(x) - clr(y) \|_2$$

For low-dose non-focal analyses, *A. caviae* was removed and the remaining 15 species were reclosed before CLR transformation.

**Community-state segmentation**

Community-state landscapes were segmented separately for each of the eight constructed landscapes with a custom greedy spatial algorithm. For a partition $\mathcal{R}$, the objective combined generalized Jensen-Shannon within-state compositional loss with the number of boundaries *B* between adjacent environmental cells:

$$J(R) = \sum_{R \in R} L_{GJS}(R) + \lambda B(R)$$

At each step, adjacent states were merged when the change in this objective was negative. The baseline boundary penalty was $\lambda$=0.15, and a merge was rejected if the resulting state exceeded a Bray-Curtis diameter of 0.40. Sensitivity analyses varied $\lambda$ and the maximum permitted Bray-Curtis diameter over the prespecified combinations shown in Figure S1. Two summaries of each segmented landscape are reported. The number of community states is the number of states in the partition. The typical state size is $\Sigma s^2/N$ over states of size *s* in a landscape of *N* treatment cells, that is, the size of the state containing a randomly chosen treatment cell. It is reported in units of treatment cells and is larger than the plain mean state size whenever states differ in size.

Landscape topology was compared between focal-species histories on the state boundaries. Each segmented landscape was coded as a binary vector over its 108 adjacent-cell edges (60 between neighbouring carbon levels and 48 between neighbouring streptomycin levels), with 1 where the edge separated two states. Exact two-group PERMANOVA on Euclidean distances between these vectors tested whether boundary configurations differed between histories; one minus the adjusted Rand index between whole partitions gave a sensitivity analysis. The boundary incidence of an environmental cell was the fraction of its adjacent edges that crossed a state boundary. Over the 64 cells, or over the 16 carbon levels after averaging boundary incidence across streptomycin, the spatial heterogeneity of the history effect was $T = \Sigma(d\_k - \bar{d})^2$, where d_k is the evolved-minus-ancestral difference in mean boundary incidence; its exact *P* value was obtained by recomputing T for all 12,870 allocations of history labels to the 16 constructed landscapes. To describe where boundaries changed, boundary incidence was averaged within low (carbon levels 1–5), intermediate (6–11) and high (12–16) carbon and compared between histories, with Holm correction across the three regions. Boundary orientation was summarized as the number of boundary edges per landscape between neighbouring carbon levels and between neighbouring streptomycin levels, compared between histories by exact allocation tests with Holm correction across the two axes.

**Conditionality of focal-species success**

We tested whether changes in community-state boundaries were associated with the environmental sensitivity of resistance-evolved *A. caviae*. For each of the 64 environments, sensitivity was calculated as the magnitude of the local gradient of its mean relative abundance across the 16-carbon × 4-streptomycin grid. Gradients were calculated in grid-index space using central differences, with one-sided differences at the edges. This gave the two environmental axes equal weight in the gradient calculation. We then calculated the evolved-minus-ancestral difference in boundary incidence at each environment (Fig. 2D) and tested its association with focal-species sensitivity using Spearman correlation, separately for three- and 16-species communities.

Significance was assessed by exact randomization of evolutionary-history labels across the 16 constructed landscapes (6,435 distinct allocations). Both the abundance gradients and boundary differences were recalculated for each allocation, preserving the spatial structure of the data within landscapes. Confidence intervals were obtained from 10,000 bootstrap resamples of constructed landscapes within each evolutionary history. Because abundance gradients can become small when a species approaches fixation, we also calculated partial Spearman correlations controlling for mean resistance-evolved *A. caviae* abundance.

**Randomization inference and multiplicity control**

Individual community cultures were the biological replicates at each environmental condition. For analyses summarizing complete response surfaces or averages across environmental conditions, the eight fixed constructed landscapes per evolutionary history served as analytical units. Each biological community therefore contributed only once to a landscape-level statistic. For comparisons between ancestral and resistance-evolved histories, the test statistic was the difference in mean values between the two groups. For within-history contrasts, it was the mean of the eight constructed-landscape differences:

$$T_{ind}=\acute{y}_E-\acute{y}_A, T_{paired}=\acute{d}$$

Between-history comparisons used two-sided exact randomization over all 12,870 possible allocations of the 16 constructed landscapes into two groups of eight. Within-history contrasts used all $2^8$=256 possible sign assignments of the eight landscape-level differences. These sign-flip tests were applied to constructed-landscape contrasts; the fixed replicate-index grouping across environmental conditions did not represent biological pairing. For two-sided scalar tests, the exact *P* value was the proportion of randomizations yielding an absolute test statistic at least as large as observed.

Dose-level Shannon Index and resistance-filtering summaries were averaged across the 16 carbon treatments within each constructed landscape before testing. Species-specific ΔCLR contrasts between evolutionary histories were tested separately at each antibiotic concentration, with Benjamini–Hochberg correction across the 16 species.

Multivariate landscape effects were tested using exact two-group PERMANOVA. Each constructed landscape was represented by either its 64 Shannon Index values or the relevant CLR response coordinates. Evolutionary-history labels were permuted among the 16 constructed landscapes. The pseudo-*F* statistic was:

$$F=\frac{SS_B/(g-1)}{SS_W/(N-g)}$$

where $SS_B$ and $SS_W$ are the between-group and within-group sums of squares, respectively, $g$=2 groups and $N$=16 constructed landscapes. The exact $P$ value was the proportion of permutations yielding a pseudo-$F$ statistic greater than or equal to the observed value. Confidence intervals were calculated as 95% percentile intervals from 10,000 bootstrap resamples of the eight constructed landscapes within each evolutionary history. Holm adjustment was applied to prespecified small families of tests, and Benjamini-Hochberg false-discovery-rate correction was used for species-wide comparisons.

**Low-dose focal/non-focal coupling**

At 2 µg ml$^{-1}$ streptomycin, the focal response within each carbon treatment was the change in *A. caviae* relative abundance from 0 to 2 µg ml$^{-1}$. For each of the 16 fixed carbon levels, the x-coordinate was the evolved-minus-ancestral difference in that focal response, with each history mean estimated from $N$=8 independent biological replicates. The y-coordinate was the Aitchison distance between the corresponding ancestral- and evolved-history non-focal ΔCLR centroids after removing and reclosing *A. caviae*. The across-carbon association was summarized by Spearman ρ. Its exact $P$ value was obtained by reallocating evolutionary-history labels among the 16 constructed landscapes and recalculating all 16 carbon-level summary points and ρ for each allocation. Uncertainty was estimated by bootstrap resampling the eight constructed landscapes within each history. Thus, the 16 carbon treatments define environmental conditions rather than biological replicates, and the across-condition grouping is the fixed analytical construction described above.

Because the history effect on the focal response at 2 µg ml$^{-1}$ changes sign along the carbon gradient, its carbon average can be small even when the response landscapes differ. The focal response was therefore also compared between histories at each of the 16 carbon levels (exact allocation tests, Benjamini-Hochberg correction across levels) and summarized by the mean absolute history difference across carbon levels and by the distance between history centroids of the 16-level response landscapes, both tested by exact allocation randomization.

**Monoculture growth traits and competitive position**

Monoculture growth traits were measured with six biological replicates per strain × carbon × streptomycin condition. For the three dominant species, endpoint competitive position $q_i$ was the CLR coordinate defined above. Kinetic-trait analyses used 0, 2 and 20 µg ml$^{-1}$ streptomycin; 200 µg ml$^{-1}$ was excluded because ancestral *A. caviae* and *P. chlororaphis* showed negligible growth, making rate and lag estimates unsuitable for competitive-trait inference. Let $r_i$ and $\ell_i$ denote maximum growth rate and lag time of species $i$, and $\bar{r}$ and $\bar{\ell}$ their means across the three species in the same environment. The centered traits were standardized, and within each of the eight constructed community landscapes per history the following ordinary least-squares model was fitted across species × carbon × dose observations:

$$q_i=\beta_0+\beta_r z\left(r_i-\acute{r}\right)+\beta_l z\left(l_i-\acute{l}\right)+\varepsilon_i$$

The coefficients $\beta r$ and $\beta \ell$ are partial slopes: each measures the association of one growth trait with competitive position while controlling for the other. For each history, the mean of the eight replicate-specific slopes was tested against zero by exact sign-flip randomization; the evolved-minus-ancestral difference in slopes was tested by exact allocation randomization of the 16 replicate-specific slope estimates. The same model was fitted separately at each dose for dose-resolved analyses. Joint model $R^2$ was decomposed between growth rate and lag using the two-predictor Shapley/LMG contribution:

$$\phi_r = \frac{1}{2}\left\{R^2(r) + \left[R^2(r,l) - R^2(l)\right]\right\}, \phi_l = R^2(r,l) - \phi_r$$

where $\varphi r$ and $\varphi \ell$ sum to the joint two-trait $R^2$. This decomposition was used descriptively to show how the relative contribution of the two traits changed with streptomycin concentration; inferential tests were applied to the replicate-specific partial slopes rather than to the Shapley fractions.

### Supplementary Figures

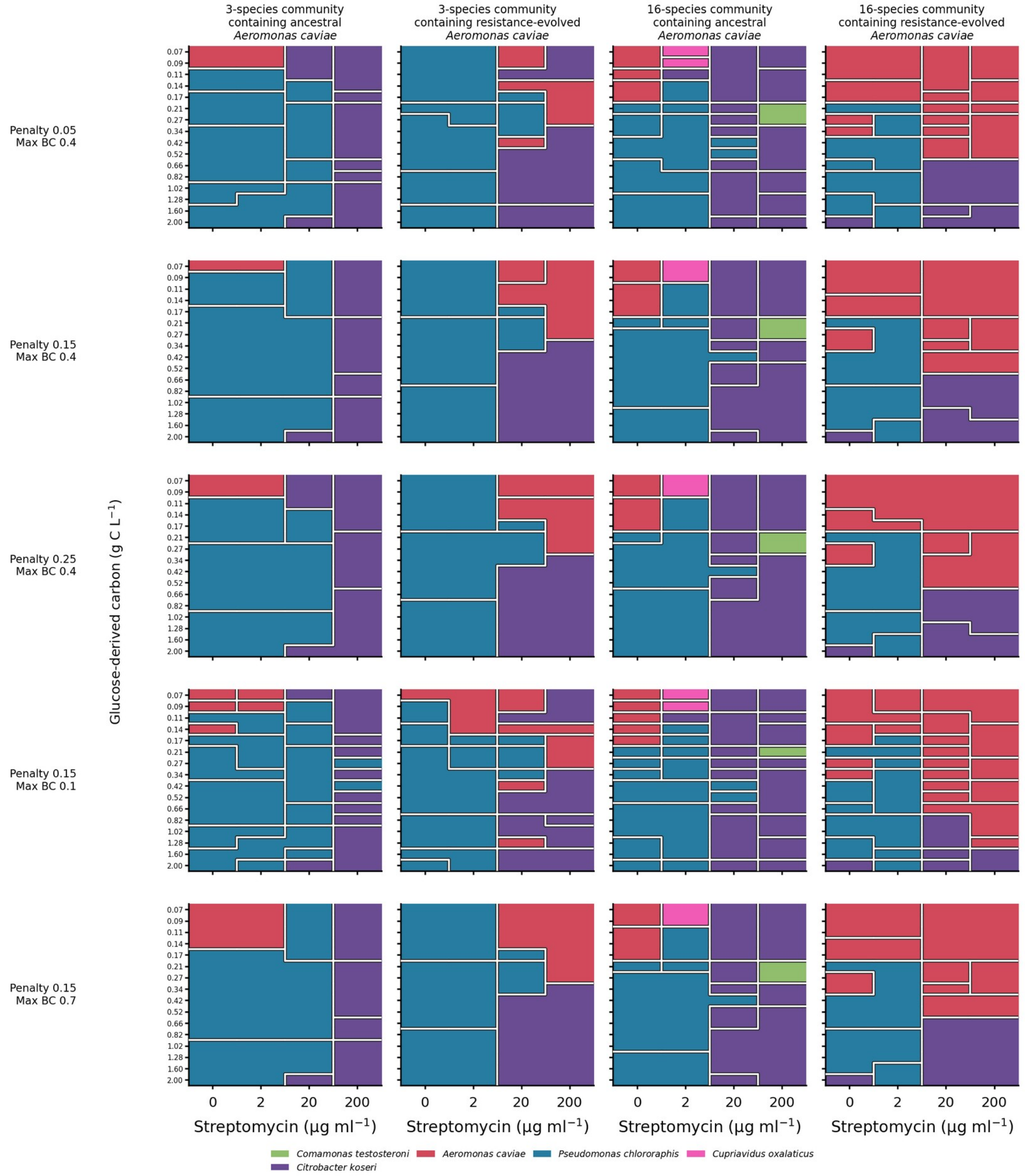


**Figure S1 |** Segmentation sensitivity across parameter choices. Mean-composition landscapes across the prespecified segmentation parameter combinations. Colour indicates the dominant species of each state centroid and white lines indicate state boundaries. Each treatment mean is based on *N*=8 replicates.

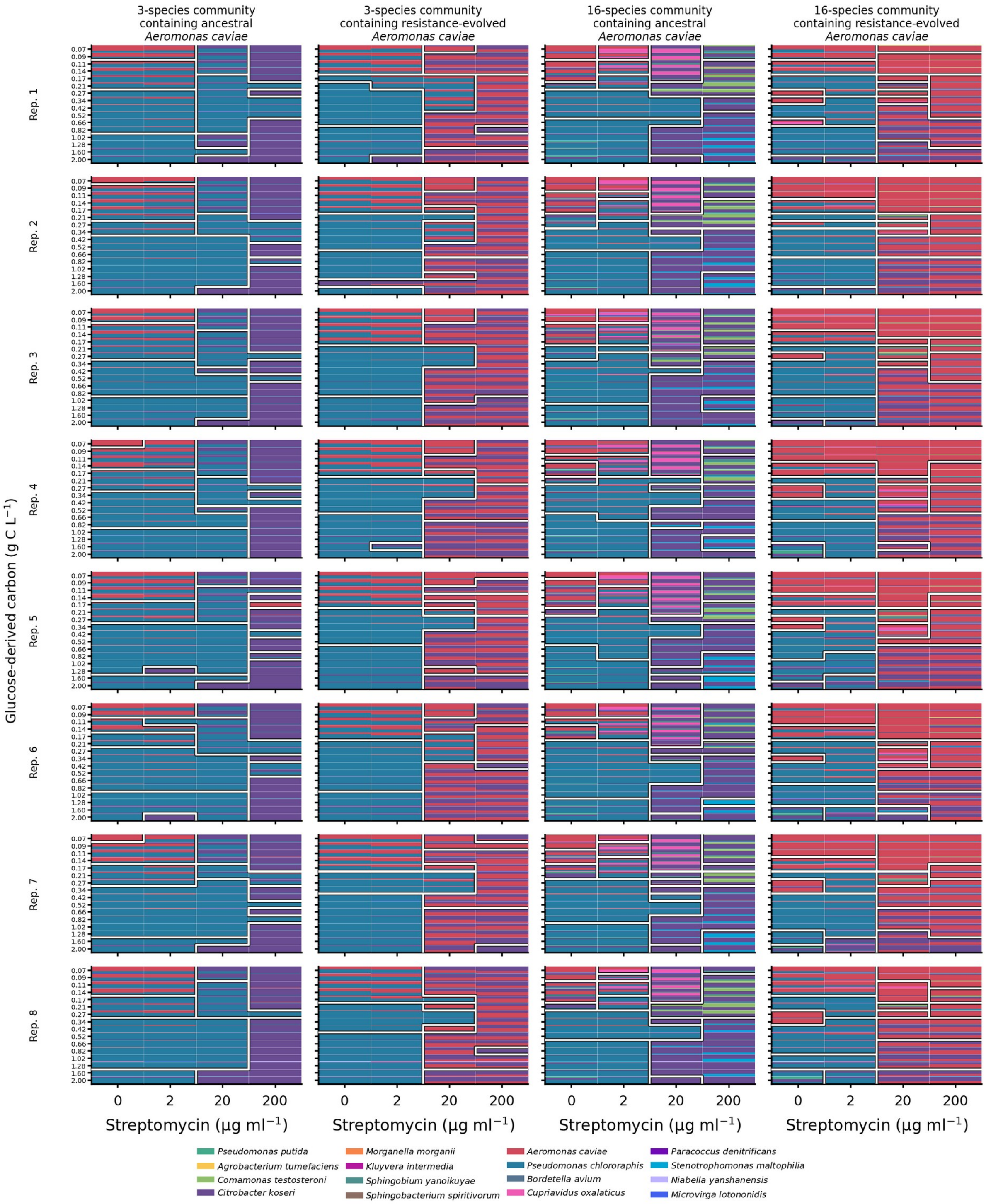


**Figure S2 |** Community composition with replicate-specific state boundaries. Each row is one biological replicate and each column is one community treatment. Every environmental cell shows untransformed species relative abundance; white lines show segmentation calculated independently for that replicate. *N*=8 replicates per treatment.

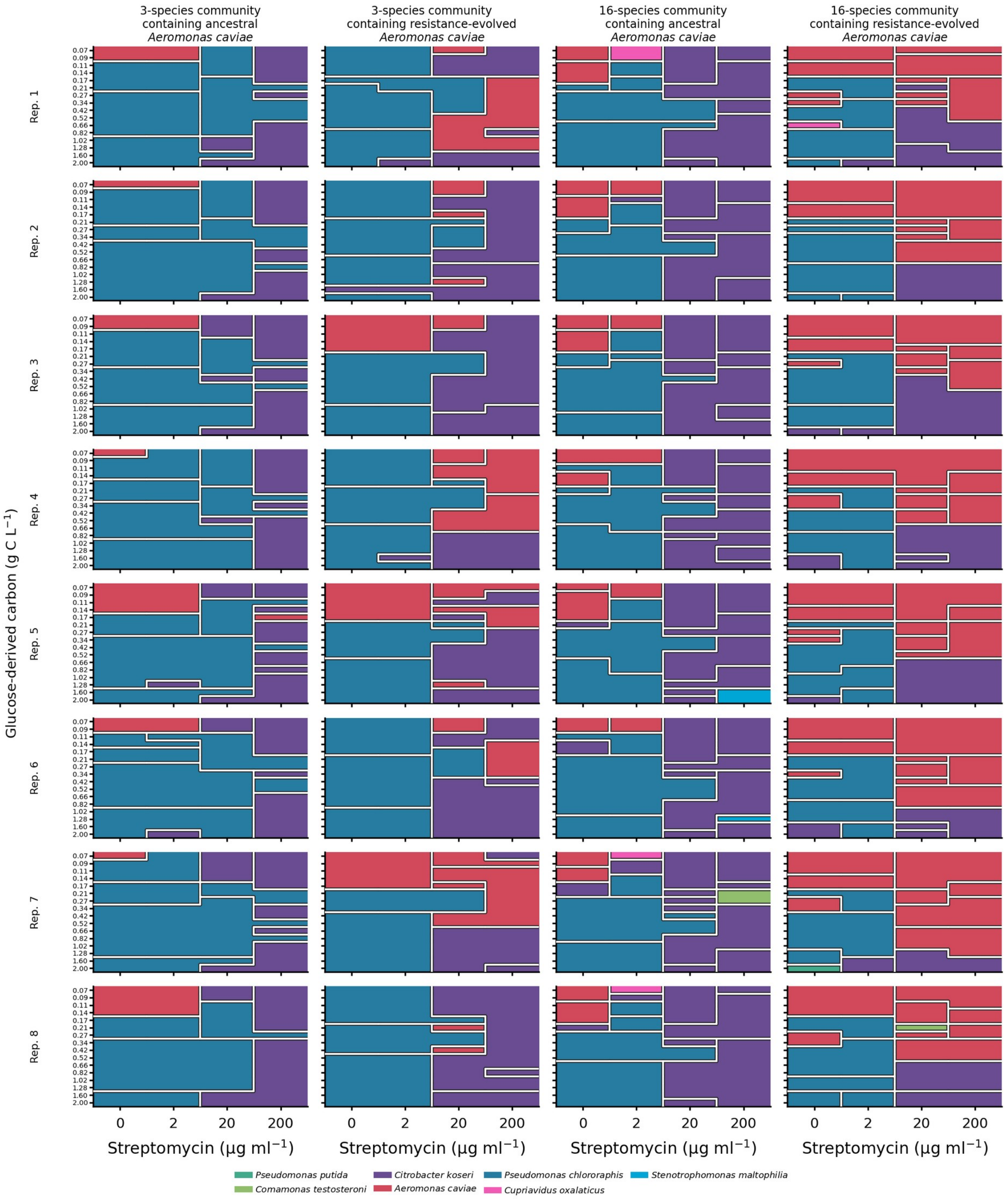


**Figure S3 |** Dominant-species state segmentations. Replicate-specific segmentations are shown with each state coloured by the dominant species of its centroid and state boundaries in white. The legend includes only species that dominate at least one segmented state. *N*=8 replicates per treatment.

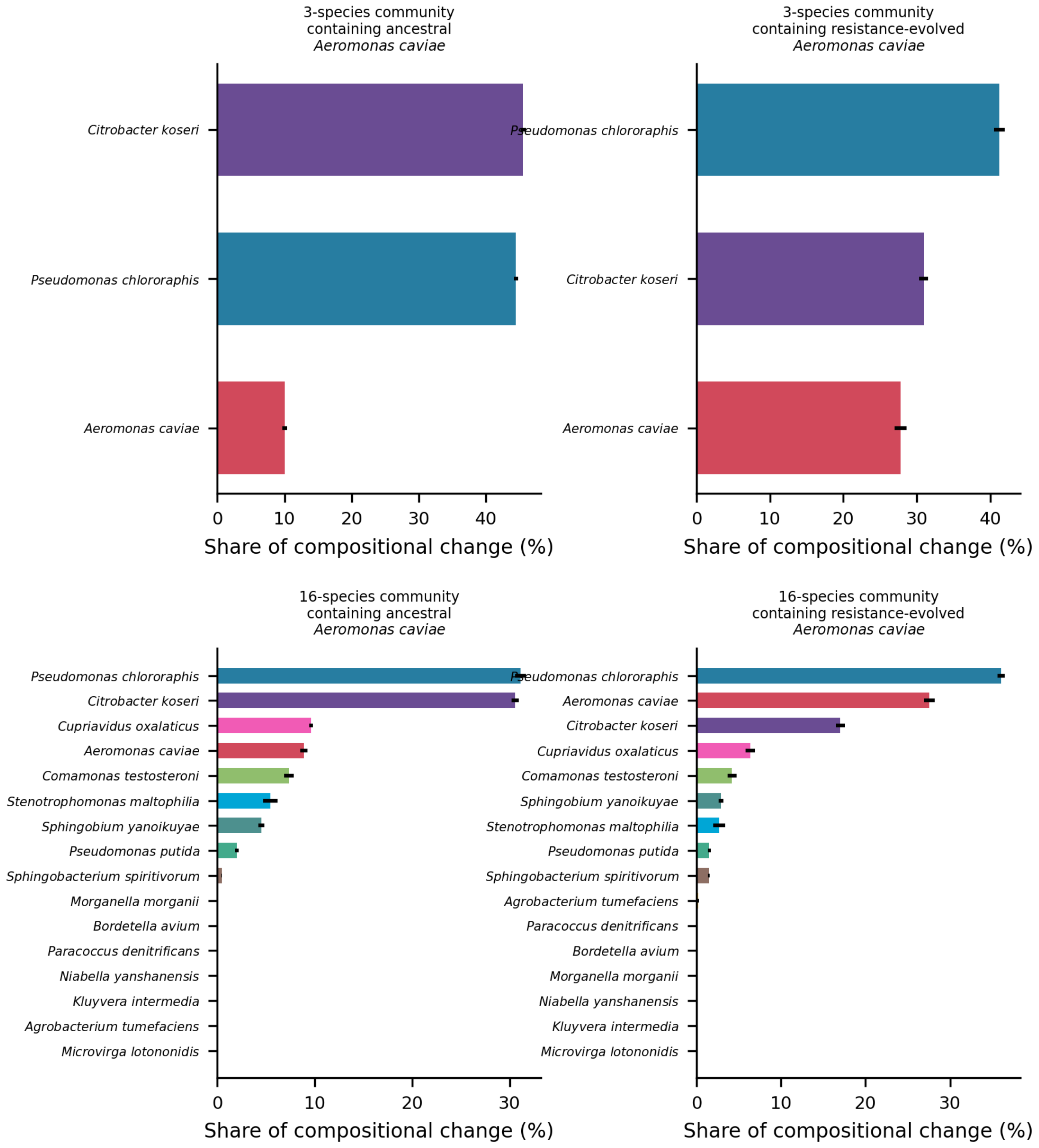


**Figure S4 |** Species contributions across segmentation boundaries. Mean share of absolute compositional change across replicate-specific segmentation boundaries attributable to each species. Error bars show s.e.m. across *N*=8 replicates per treatment.

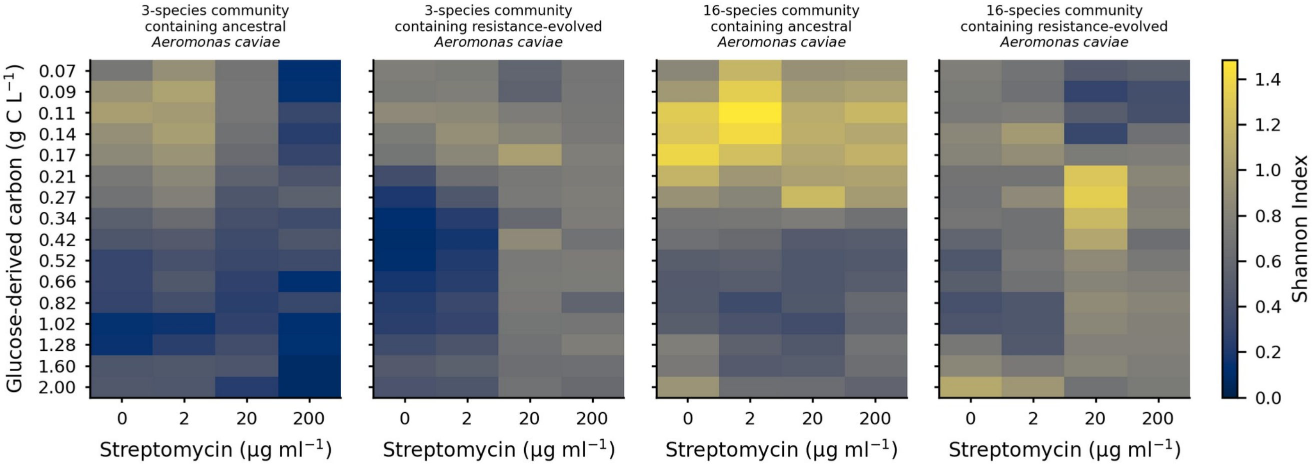


**Figure S5 |** Absolute alpha-diversity landscapes. Mean Shannon Index at each carbon-by-streptomycin condition for all four experimental communities. Each condition mean is based on *N*=8 replicates.

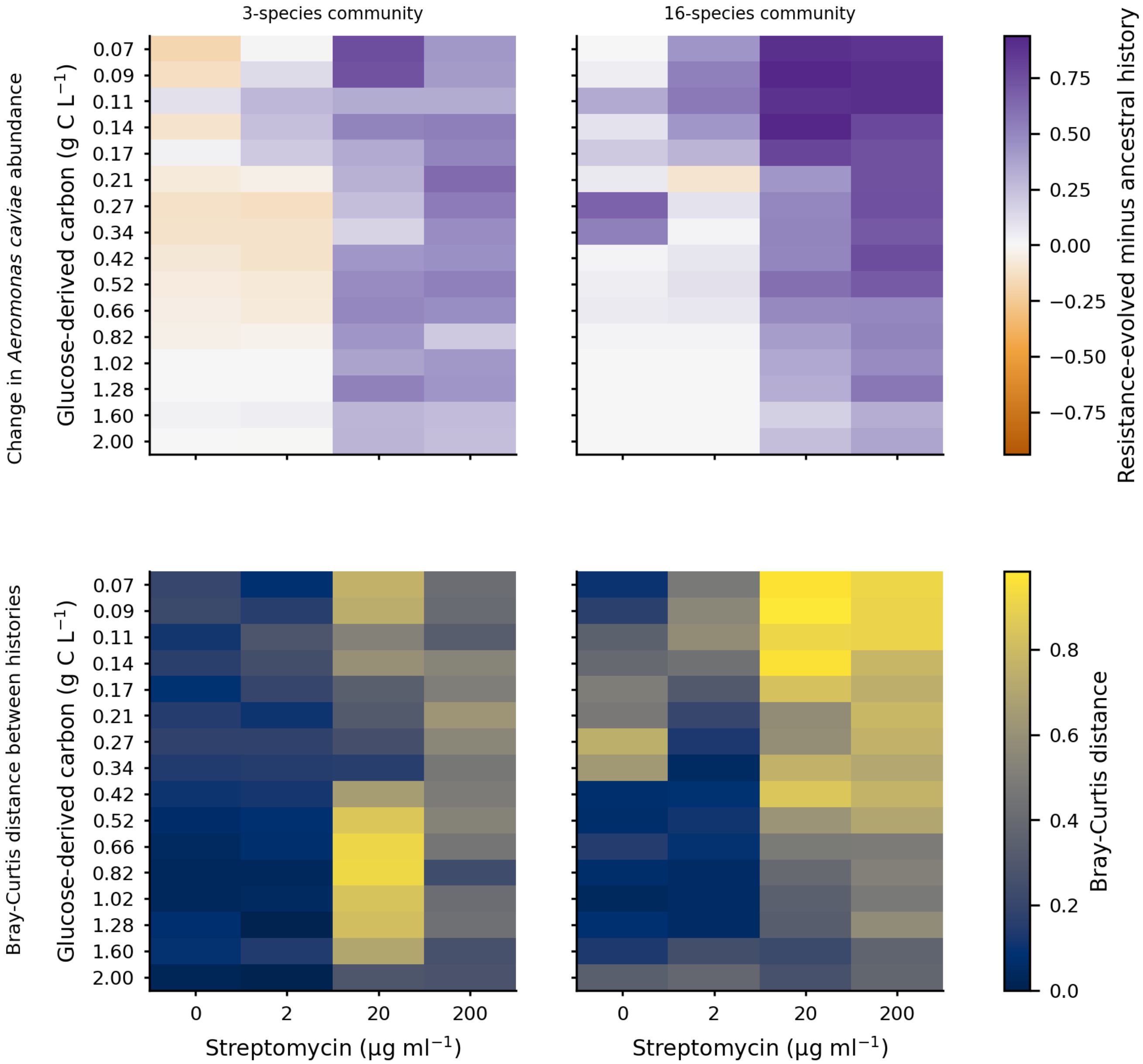


**Figure S6 |** Condition-level effects of focal-species evolutionary history. Upper row, evolved-minus-ancestral change in *A. caviae* relative abundance. Lower row, Bray-Curtis dissimilarity between ancestral-history and resistance-evolved-history mean compositions at each condition. Each history mean is based on *N*=8 replicates.

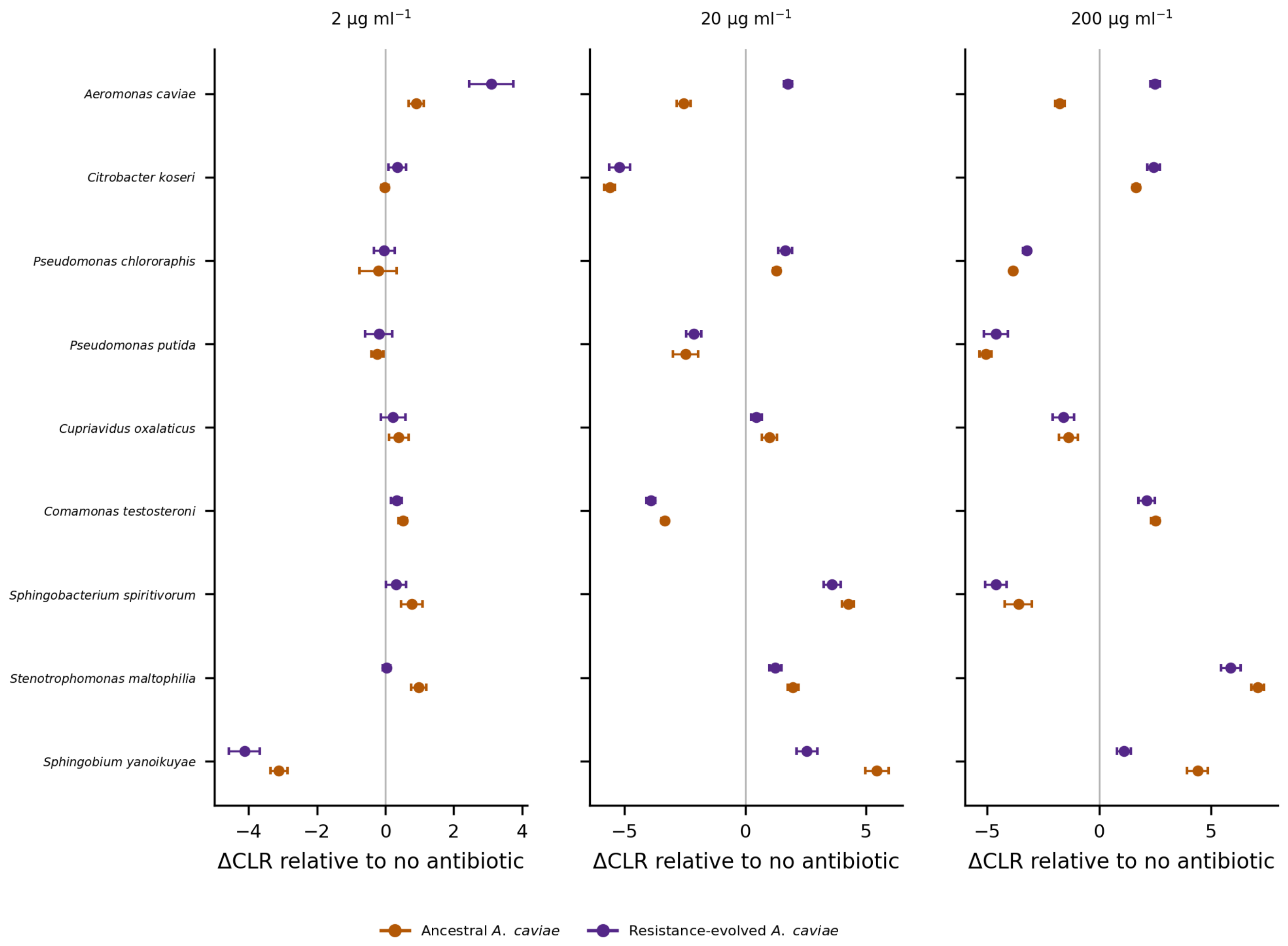


**Figure S7 |** Species-specific antibiotic responses across doses. ΔCLR is the change in a species' centred-log-ratio abundance from the matched no-antibiotic condition at the same carbon concentration; positive values indicate an increase relative to the geometric mean abundance of all species and negative values a decrease. Points show carbon-averaged means and error bars show 95% confidence intervals across $N$=8 replicates per history.

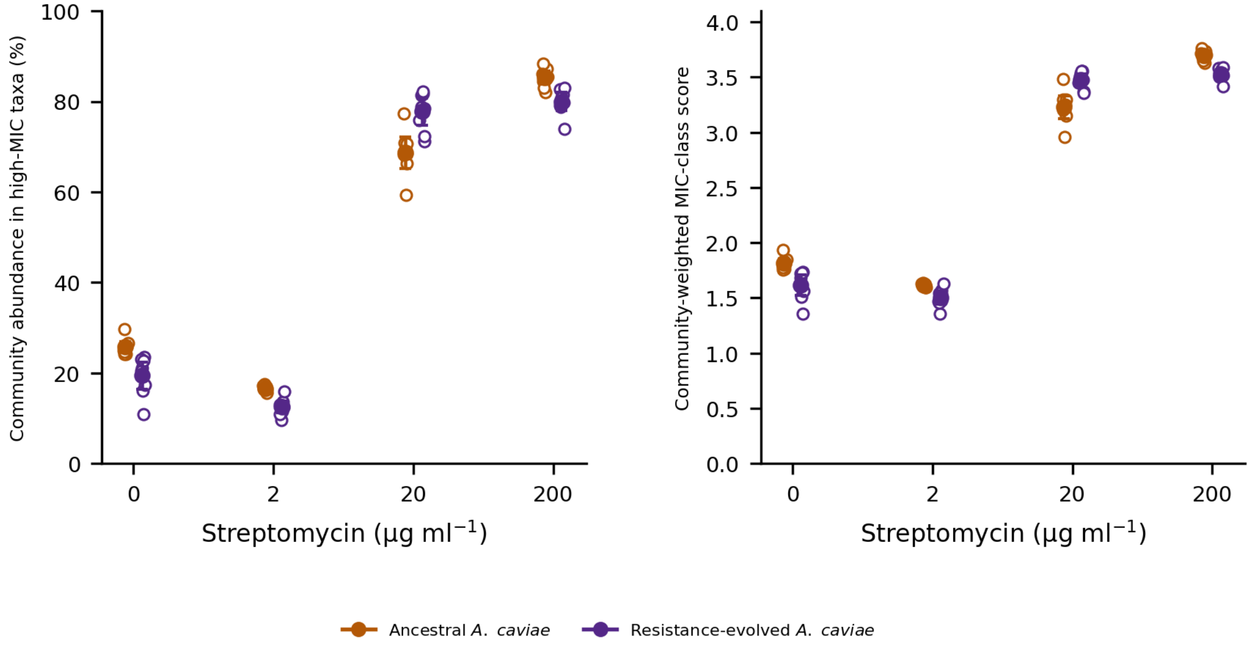


**Figure S8 |** Community-level resistance enrichment. Left, percentage of non-focal abundance carried by high-MIC taxa; right, community-weighted MIC-class score, both excluding focal *A. caviae*. Open points are biological replicates; filled points and error bars show means ± 95% confidence intervals across *N*=8 replicates per history.

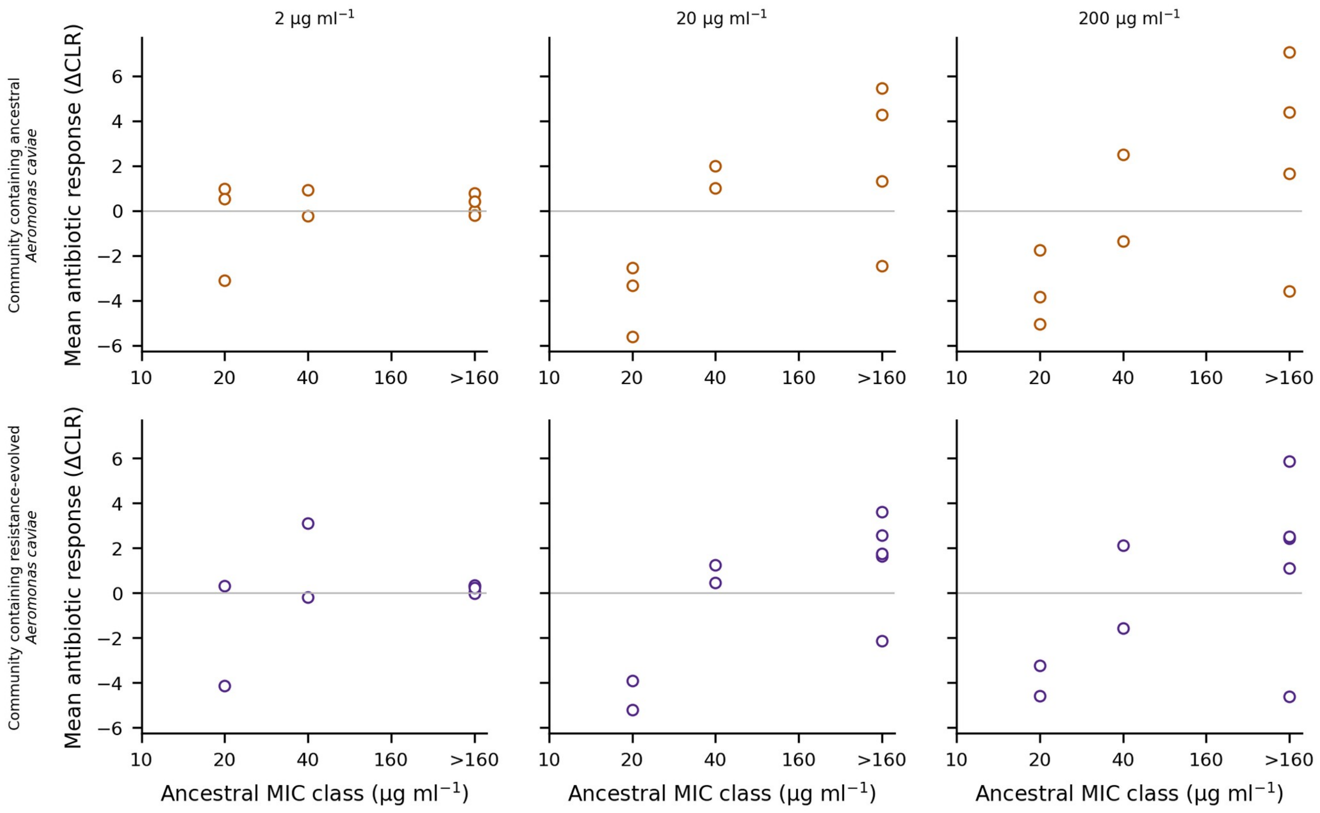


**Figure S9 |** Species resistance class and antibiotic-induced response. Species-level mean ΔCLR response is shown against ancestral streptomycin MIC class at each non-zero dose for communities containing ancestral or resistance-evolved *A. caviae*. Each point is a species mean calculated from *N*=8 replicates; permutation statistics are reported in Supplementary Table S11.

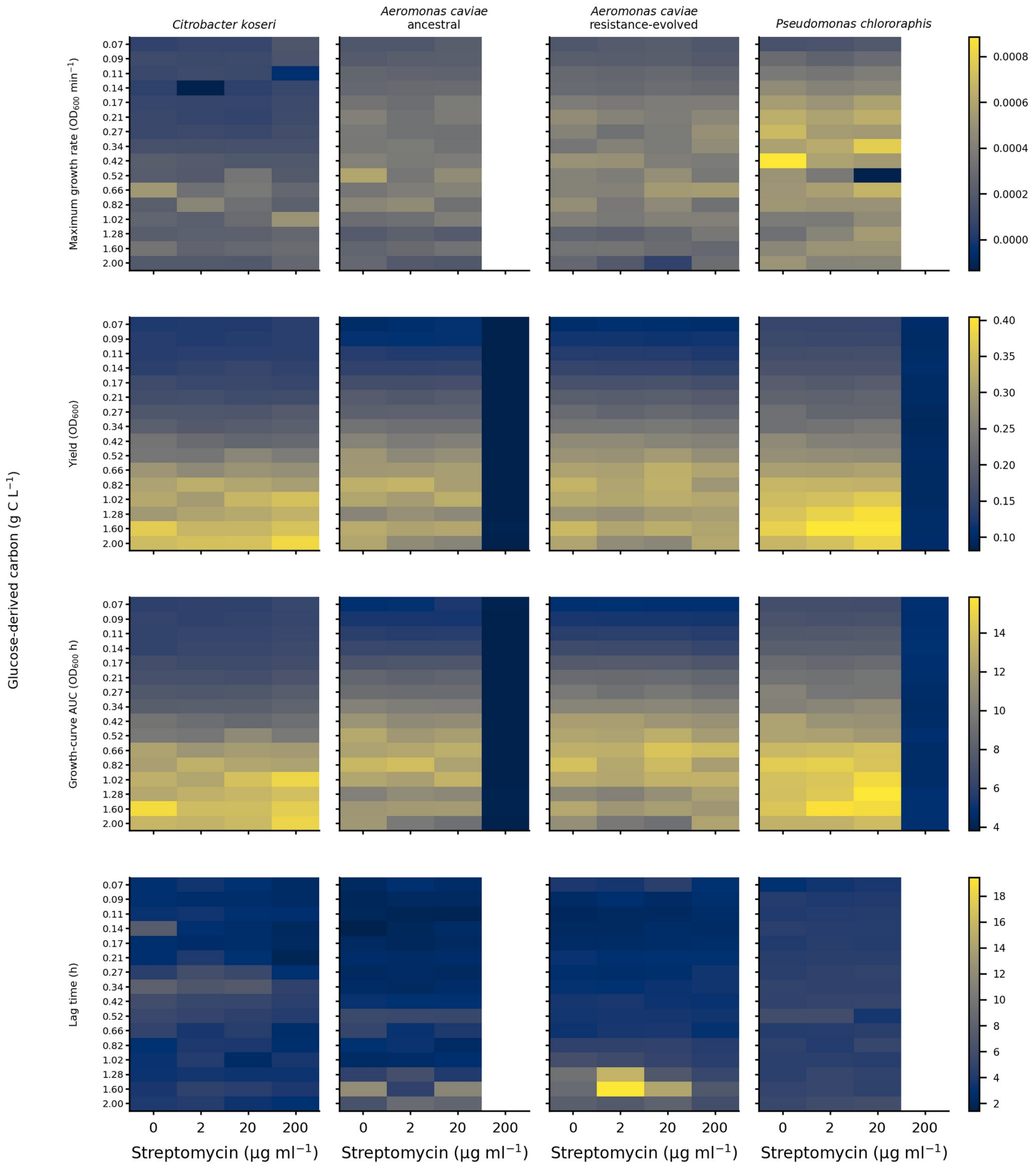


**Figure S10 |** Monoculture growth traits across the resource-antibiotic landscape. Mean maximum growth rate, yield, growth-curve AUC and lag time are shown across 16 carbon and four streptomycin levels for *C. koseri*, ancestral and resistance-evolved *A. caviae*, and *P. chlororaphis*. Means use *N*=6 monoculture replicates per strain-condition. Maximum-growth-rate and lag estimates are omitted at 200 µg ml$^{-1}$ for ancestral *A. caviae* and *P. chlororaphis* because growth was negligible.

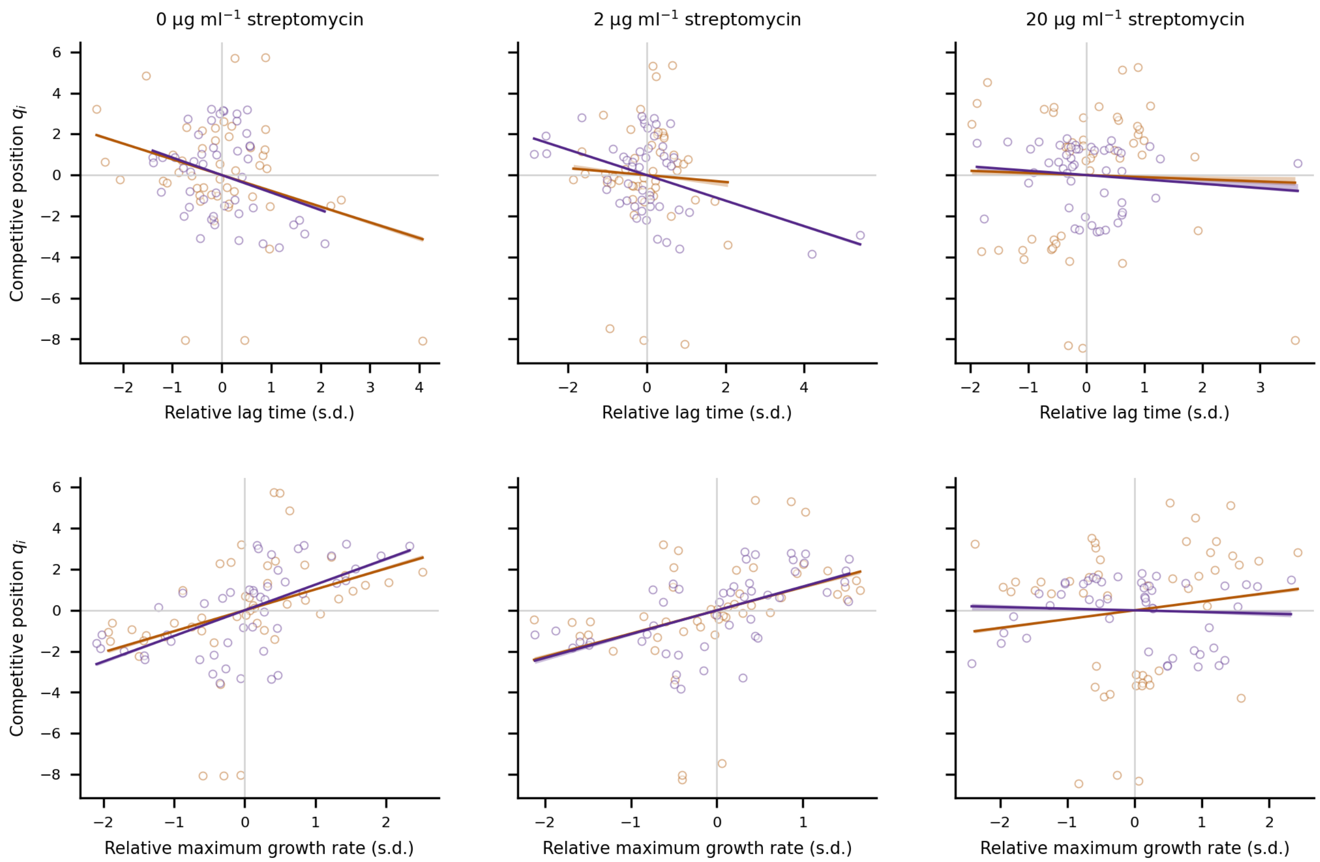


**Figure S11 |** Dose-resolved associations of monoculture growth traits with competitive position. Rows show relative lag time and relative maximum growth rate; columns show 0, 2 and 20 µg ml$^{-1}$ streptomycin. Points are species × carbon condition means based on $N$=6 monoculture and $N$=8 community replicates. Lines show median partial slopes from the joint rate-plus-lag model and shading shows bootstrap 95% confidence intervals of the median obtained by resampling $N$=8 community replicates. Slopes and Holm-adjusted exact sign-flip $P$ values are given in Supplementary Table S13 and joint $R^2$ in Supplementary Table S14.

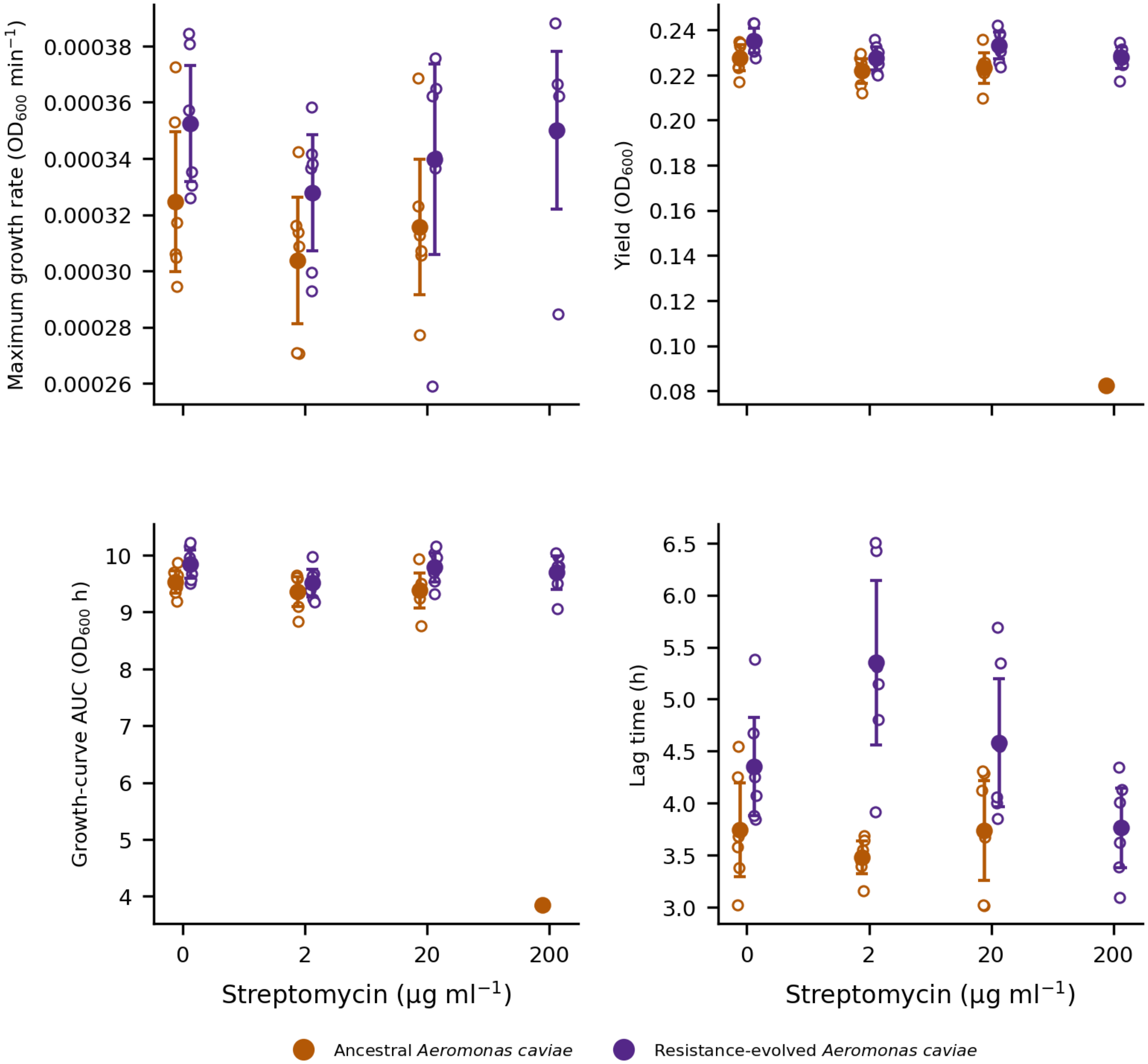


**Figure S12 |** Growth phenotype of ancestral and resistance-evolved *A. caviae*. Open points are biological monoculture replicates after averaging across the 16 carbon levels within replicate and antibiotic dose; filled points and error bars show means ± 95% confidence intervals (*N*=6). Maximum-growth-rate and lag estimates are omitted for ancestral *A. caviae* at 200 μg $ml^{-1}$ because growth was negligible.

### Supplementary Tables

**Supplementary Table S1. Primary tests of community-state landscape structure.** Boundary configuration was compared between histories by exact two-group PERMANOVA on the 108-edge boundary vectors of the 16 constructed landscapes (sensitivity analysis: 1 − adjusted Rand distance between partitions; all 12,870 allocations). Heterogeneity tests ask whether the evolved-minus-ancestral change in boundary incidence varies across the 64 environments or along the 16 carbon levels. Regional boundary incidence and state metrics are evolved-versus-ancestral contrasts tested by exact allocation randomization (*N*=8 per history) with 10,000-replicate bootstrap confidence intervals; Holm adjustment is across the three carbon regions, the two boundary axes and the three state metrics, each family separately within each richness.

| Richness | Metric or test | Ancestral | Evolved | Δ history or statistic | 95% CI or $R^2$ | *P* exact | *P* Holm |
|---|---|---|---|---|---|---|---|
| 3 | Boundary configuration (PERMANOVA) | — | — | F = 6.58 | $R^2$ = 0.320 | 0.000155 | — |
| 3 | Partition (1 − adjusted Rand PERMANOVA) | — | — | F = 11.04 | $R^2$ = 0.441 | 0.000155 | — |
| 3 | Boundary sharing (mean Jaccard) | — | — | within 0.50, between 0.28 | — | — | — |
| 3 | Heterogeneity of boundary change (64 environments) | — | — | — | — | 0.000155 | — |
| 3 | Heterogeneity of boundary change (16 carbon levels) | — | — | — | — | 0.0073 | — |
| 3 | Boundary incidence, low carbon (0.07–0.17 g C $l^{-1}$) | 0.422 | 0.324 | -0.098 | [-0.150, -0.051] | 0.00218 | 0.00653 |
| 3 | Boundary incidence, intermediate carbon (0.21–0.66 g C $l^{-1}$) | 0.344 | 0.302 | -0.042 | [-0.088, -0.002] | 0.108 | 0.217 |
| 3 | Boundary incidence, high carbon (0.82–2.00 g C $l^{-1}$) | 0.293 | 0.271 | -0.022 | [-0.080, 0.035] | 0.488 | 0.488 |
| 3 | Boundaries along the carbon axis (per landscape) | 15.12 | 13.25 | -1.88 | [-4.25, 0.50] | 0.21 | 0.21 |
| 3 | Boundaries along the streptomycin axis (per landscape) | 23.38 | 20.75 | -2.62 | [-4.25, -0.88] | 0.0216 | 0.0432 |
| 3 | Number of states | 11.00 | 9.00 | -2.000 | [-3.625, -0.5] | 0.0502 | 0.151 |
| 3 | Typical state size (cells) | 9.98 | 11.082 | 1.102 | [-0.574, 2.906] | 0.266 | 0.532 |
| 3 | Largest-state fraction | 0.277 | 0.250 | -0.027 | [-0.086, 0.033] | 0.449 | 0.532 |
| 16 | Boundary configuration (PERMANOVA) | — | — | F = 6.18 | $R^2$ = 0.306 | 0.000155 | — |
| 16 | Partition (1 − adjusted Rand PERMANOVA) | — | — | F = 10.47 | $R^2$ = 0.428 | 0.000155 | — |
| 16 | Boundary sharing (mean Jaccard) | — | — | within 0.55, between 0.35 | — | — | — |
| 16 | Heterogeneity of boundary change (64 environments) | — | — | — | — | 0.000155 | — |
| 16 | Heterogeneity of boundary change (16 carbon levels) | — | — | — | — | 0.000155 | — |
| 16 | Boundary incidence, low carbon (0.07–0.17 g C $l^{-1}$) | 0.583 | 0.311 | -0.272 | [-0.320, -0.221] | 0.000155 | 0.000466 |
| 16 | Boundary incidence, intermediate carbon (0.21–0.66 g C $l^{-1}$) | 0.405 | 0.491 | 0.086 | [0.043, 0.132] | 0.00171 | 0.00342 |
| 16 | Boundary incidence, high carbon (0.82–2.00 g C $l^{-1}$) | 0.322 | 0.368 | 0.045 | [-0.021, 0.114] | 0.245 | 0.245 |

| Richness | Metric or test | Ancestral | Evolved | Δ history or statistic | 95% CI or $R^2$ | *P* exact | *P* Holm |
|---|---|---|---|---|---|---|---|
| 16 | Boundaries along the carbon axis (per landscape) | 16.75 | 19.62 | 2.88 | [1.12, 4.62] | 0.0141 | 0.0141 |
| 16 | Boundaries along the streptomycin axis (per landscape) | 31.12 | 24.75 | -6.38 | [-8.50, -4.25] | 0.000155 | 0.000311 |
| 16 | Number of states | 15.00 | 15.38 | 0.375 | [-1.25, 1.875] | 0.72 | 0.72 |
| 16 | Typical state size (cells) | 7.902 | 6.730 | -1.172 | [-2.016, -0.293] | 0.0289 | 0.0578 |
| 16 | Largest-state fraction | 0.219 | 0.176 | -0.043 | [-0.074, -0.016] | 0.0191 | 0.0573 |

**Supplementary Table S2. Dose-level alpha-diversity contrasts.** Shannon Index was first averaged across the 16 carbon treatments within each constructed landscape. Evolved-versus-ancestral differences were tested by exact allocation randomization (*N*=8 constructed landscapes per history); Holm adjustment is across the four streptomycin doses within each richness.

| Richness | Strep. (μg ml$^{-1}$) | Δ Shannon Index | 95% CI | *P* exact | *P* Holm |
|---|---|---|---|---|---|
| 3 | 0 | -0.141 | [-0.155, -0.126] | 0.000155 | 0.000622 |
| 3 | 2 | -0.128 | [-0.144, -0.111] | 0.000155 | 0.000622 |
| 3 | 20 | 0.261 | [0.239, 0.282] | 0.000155 | 0.000622 |
| 3 | 200 | 0.450 | [0.426, 0.475] | 0.000155 | 0.000622 |
| 16 | 0 | -0.168 | [-0.196, -0.139] | 0.000155 | 0.000622 |
| 16 | 2 | -0.122 | [-0.143, -0.101] | 0.000155 | 0.000622 |
| 16 | 20 | 0.054 | [0.024, 0.083] | 0.00559 | 0.00559 |
| 16 | 200 | -0.091 | [-0.120, -0.058] | 0.000622 | 0.00124 |

**Supplementary Table S3. Global effect of focal-species history on the alpha-diversity landscape.** Each constructed landscape was represented by its 64-dimensional Shannon Index surface. Exact two-group PERMANOVA tested the history effect over all 12,870 allocations of eight labels among 16 constructed landscapes.

| Richness | Model | Pseudo-*F* | $R^2$ | *P* exact | Allocations |
|---|---|---|---|---|---|
| 3 | Exact PERMANOVA | 59.05 | 0.808 | 0.000155 | 12870 |
| 16 | Exact PERMANOVA | 37.22 | 0.727 | 0.000155 | 12870 |

**Supplementary Table S4. Whole-community compositional antibiotic-response models.** At each non-zero dose, each biological replicate was represented by its carbon-averaged species-wise ΔCLR vector. Exact PERMANOVA in Euclidean CLR (Aitchison) space tested the effect of focal-species history (*N*=8 landscapes per history).

| Richness | Strep. (µg $ml^{-1}$) | Pseudo-*F* | $R^2$ | *P* exact |
|---|---|---|---|---|
| 3 | 2 | 4.632 | 0.249 | 0.00109 |
| 3 | 20 | 1033 | 0.987 | 0.000155 |
| 3 | 200 | 325.7 | 0.959 | 0.000155 |
| 16 | 2 | 14.47 | 0.508 | 0.000155 |
| 16 | 20 | 64.85 | 0.822 | 0.000155 |
| 16 | 200 | 53.79 | 0.793 | 0.000155 |

**Supplementary Table S5. Direct focal-species response at 2 μg $ml^{-1}$ streptomycin.** The response is the change in *A. caviae* relative abundance from 0 to 2 μg $ml^{-1}$, averaged across the 16 carbon treatments within each constructed landscape. Histories were compared by exact allocation randomization; Holm adjustment is across the two community richness levels. Because the history effect changes sign along the carbon gradient (Fig. 4A,B), the response landscapes were also compared: the mean absolute history difference across carbon levels was 0.080 (three species) and 0.186 (16 species), and the distance between history centroids of the 16-level response landscapes 0.54 and 1.11 (exact $P$=0.00016 for all four tests).

| Richness | Ancestral Δ abundance | Evolved Δ abundance | Δ history | 95% CI | *P* exact | *P* Holm |
|---|---|---|---|---|---|---|
| 3 | -0.018 | 0.052 | 0.069 | [0.059, 0.080] | 0.000155 | 0.000311 |
| 16 | -0.072 | -0.042 | 0.029 | [0.006, 0.053] | 0.0375 | 0.0375 |

**Supplementary Table S6. Low-dose coupling between focal and non-focal responses.** The association across carbon treatments was evaluated with Spearman correlation; inference permuted history labels across the 16 constructed landscapes and confidence intervals bootstrap-resampled the eight constructed landscapes within each history. Non-focal landscape divergence was tested by exact allocation randomization.

| Analysis | Statistic | Estimate | 95% CI | *P* exact | *N* |
|---|---|---|---|---|---|
| Focal-response history effect vs non-focal divergence | Spearman ρ | 0.706 | [0.450, 0.797] | 0.0113 | 8/history; 16 carbon |
| Non-focal response landscape history effect | RMS Aitchison separation | 9.09 | — | 0.000155 | 8/history |

**Supplementary Table S7. Log-ratio sensitivity analysis: species-specific history effects on the 2 μg $ml^{-1}$ antibiotic response.** Main-text species-level contrasts are reported as changes in relative abundance (Figure 4d); this table gives the log-ratio version for comparison. For each non-focal species retained in the CLR basis, the carbon-averaged ΔCLR response was compared between histories by exact allocation randomization of *N*=8 constructed landscapes per history. *q* values are Benjamini-Hochberg adjusted across the nine species in that basis, which includes focal *A. caviae*. The focal species is not listed here because its own log-ratio coordinate is governed by the zero-replacement constant (Supplementary Methods).

| Species | ΔCLR history effect | 95% CI | *P* exact | *q* BH |
|---|---|---|---|---|
| *Pseudomonas putida* HAMBI 6 | -1.008 | [-1.487, -0.514] | 0.00233 | 0.00699 |
| *Comamonas testosteroni* HAMBI 403 | 0.042 | [-0.349, 0.450] | 0.856 | 0.856 |
| *Citrobacter koseri* HAMBI 1287 | 0.357 | [0.093, 0.610] | 0.0233 | 0.0524 |
| *Sphingobium yanoikuyae* HAMBI 1842 | -0.461 | [-0.861, -0.054] | 0.0563 | 0.101 |
| *Sphingobacterium spiritivorum* HAMBI 1896 | 0.179 | [-0.411, 0.754] | 0.578 | 0.65 |
| *Pseudomonas chlororaphis* HAMBI 1977 | -0.188 | [-0.370, -0.006] | 0.0842 | 0.126 |
| *Cupriavidus oxalaticus* HAMBI 2164 | 2.183 | [1.524, 2.796] | 0.000155 | 0.000699 |
| *Stenotrophomonas maltophilia* HAMBI 2659 | -0.168 | [-0.615, 0.254] | 0.495 | 0.637 |

**Supplementary Table S8. Focal-species response at 2 μg ml$^{-1}$ streptomycin at each carbon level.** The response is the change in *A. caviae* relative abundance from 0 to 2 μg ml$^{-1}$ streptomycin; values are means across *N*=8 constructed landscapes per history. Histories were compared at each carbon level by exact allocation randomization (12,870 allocations), with Benjamini-Hochberg correction across the 16 carbon levels within each richness (*q* BH). These tests underlie Fig. 4A,B.

| Richness | Carbon (g C l$^{-1}$) | Ancestral Δ abundance | Evolved Δ abundance | Δ history | *P* exact | *q* BH |
|---|---|---|---|---|---|---|
| 3 | 0.07 | -0.128 | 0.064 | 0.192 | 0.000932 | 0.00213 |
| 3 | 0.09 | -0.111 | 0.142 | 0.254 | 0.000155 | 0.000497 |
| 3 | 0.11 | -0.089 | 0.096 | 0.185 | 0.000155 | 0.000497 |
| 3 | 0.14 | -0.199 | 0.155 | 0.354 | 0.000155 | 0.000497 |
| 3 | 0.17 | -0.032 | 0.138 | 0.170 | 0.00917 | 0.0183 |
| 3 | 0.21 | 0.053 | 0.081 | 0.028 | 0.0639 | 0.0929 |
| 3 | 0.27 | 0.091 | 0.077 | -0.014 | 0.454 | 0.605 |
| 3 | 0.34 | 0.025 | 0.021 | -0.003 | 0.811 | 0.879 |
| 3 | 0.42 | 0.040 | 0.009 | -0.031 | 0.000311 | 0.000829 |
| 3 | 0.52 | 0.027 | 0.015 | -0.013 | 0.046 | 0.0736 |
| 3 | 0.66 | 0.038 | 0.013 | -0.024 | 0.000155 | 0.000497 |
| 3 | 0.82 | $1.7\times10^{-4}$ | 0.001 | 0.001 | 0.824 | 0.879 |
| 3 | 1.02 | $2.3\times10^{-4}$ | $3.7\times10^{-4}$ | $1.4\times10^{-4}$ | 0.677 | 0.833 |
| 3 | 1.28 | 0.000 | -0.001 | -0.001 | 0.000155 | 0.000497 |
| 3 | 1.60 | $2.4\times10^{-5}$ | 0.013 | 0.013 | 0.937 | 0.937 |
| 3 | 2.00 | 0.001 | $-1.4\times10^{-4}$ | -0.001 | 0.0224 | 0.0398 |
| 16 | 0.07 | -0.387 | 0.048 | 0.435 | 0.000155 | 0.000311 |
| 16 | 0.09 | -0.420 | 0.068 | 0.488 | 0.000155 | 0.000311 |
| 16 | 0.11 | -0.222 | 0.005 | 0.227 | 0.000466 | 0.000829 |
| 16 | 0.14 | -0.310 | 0.027 | 0.338 | 0.000155 | 0.000311 |
| 16 | 0.17 | -0.200 | -0.103 | 0.096 | 0.019 | 0.0303 |
| 16 | 0.21 | 0.102 | -0.057 | -0.159 | 0.000155 | 0.000311 |
| 16 | 0.27 | 0.080 | -0.497 | -0.576 | 0.000155 | 0.000311 |
| 16 | 0.34 | 0.088 | -0.428 | -0.516 | 0.000155 | 0.000311 |
| 16 | 0.42 | 0.007 | 0.069 | 0.062 | 0.032 | 0.0466 |
| 16 | 0.52 | 0.043 | 0.100 | 0.058 | 0.000155 | 0.000311 |
| 16 | 0.66 | 0.063 | 0.081 | 0.018 | 0.131 | 0.174 |
| 16 | 0.82 | 0.011 | 0.010 | -0.001 | 0.844 | 0.9 |
| 16 | 1.02 | $3.8\times10^{-5}$ | $2.0\times10^{-4}$ | $1.7\times10^{-4}$ | 0.658 | 0.752 |
| 16 | 1.28 | $4.0\times10^{-5}$ | -0.002 | -0.002 | 0.000155 | 0.000311 |
| 16 | 1.60 | $-3.5\times10^{-5}$ | $-3.8\times10^{-4}$ | $-3.5\times10^{-4}$ | 0.404 | 0.498 |
| 16 | 2.00 | $-1.6\times10^{-4}$ | $-1.3\times10^{-4}$ | $2.5\times10^{-5}$ | 0.976 | 0.976 |

**Supplementary Table S9. Species-specific history effects on the 2 µg ml$^{-1}$ antibiotic response.** Values are changes in relative abundance from 0 to 2 µg ml$^{-1}$ streptomycin, averaged across the 16 carbon treatments within each constructed landscape, and the evolved-minus-ancestral difference (Δ history) with its bootstrap 95% CI. *A. caviae* (†) is measured in the whole community and the other species within the non-focal subcomposition, as in Fig. 4D. Histories were compared by exact allocation randomization (*N*=8 constructed landscapes per history). Adjusted *P* is the Benjamini-Hochberg *q* across the 15 non-focal species; for *A. caviae* (‡) it is the Holm-adjusted *P* of the primary focal-response test, which is the same contrast (Supplementary Table S5). Asterisks in Fig. 4D mark adjusted *P*<0.05.

| Species | Ancestral Δ abundance | Evolved Δ abundance | Δ history | 95% CI | *P* exact | Adjusted *P* |
|---|---|---|---|---|---|---|
| *Pseudomonas putida* HAMBI 6 | -0.086 | -0.032 | 0.053 | [0.045, 0.062] | 0.000155 | 0.00117 |
| *Agrobacterium tumefaciens* HAMBI 105 | $1.9\times10^{-8}$ | $-4.4\times10^{-5}$ | $-4.4\times10^{-5}$ | [$-1.5\times10^{-4}$, $7.1\times10^{-5}$] | 0.529 | 0.722 |
| *Comamonas testosteroni* HAMBI 403 | -0.001 | $-4.4\times10^{-4}$ | $3.3\times10^{-4}$ | [$-7.5\times10^{-5}$, 0.001] | 0.151 | 0.282 |
| *Citrobacter koseri* HAMBI 1287 | -0.090 | -0.044 | 0.045 | [0.018, 0.072] | 0.00948 | 0.0284 |
| *Morganella morganii* HAMBI 1292 | $2.2\times10^{-4}$ | $2.3\times10^{-4}$ | $1.0\times10^{-5}$ | [$-6.7\times10^{-5}$, $9.4\times10^{-5}$] | 0.816 | 0.942 |
| *Kluyvera intermedia* HAMBI 1299 | 0.000 | $2.4\times10^{-6}$ | $2.4\times10^{-6}$ | [0.000, $7.1\times10^{-6}$] | 1 | 1 |
| *Sphingobium yanoikuyae* HAMBI 1842 | 0.005 | $-1.2\times10^{-4}$ | -0.005 | [-0.006, -0.004] | 0.000155 | 0.00117 |
| *Sphingobacterium spiritivorum* HAMBI 1896 | -0.007 | -0.025 | -0.018 | [-0.029, -0.005] | 0.0204 | 0.046 |
| *Aeromonas caviae* HAMBI 1972 † | -0.072 | -0.042 | 0.029 | [0.006, 0.053] | 0.0375 | 0.0375 ‡ |
| *Pseudomonas chlororaphis* HAMBI 1977 | 0.102 | 0.017 | -0.085 | [-0.127, -0.042] | 0.0028 | 0.0105 |
| *Bordetella avium* HAMBI 2160 | -0.002 | $-2.4\times10^{-4}$ | 0.002 | [$4.7\times10^{-4}$, 0.003] | 0.0214 | 0.046 |
| *Cupriavidus oxalaticus* HAMBI 2164 | 0.078 | 0.091 | 0.013 | [-0.012, 0.032] | 0.313 | 0.522 |
| *Paracoccus denitrificans* HAMBI 2443 | $-1.9\times10^{-4}$ | -0.006 | -0.006 | [-0.009, -0.003] | 0.0014 | 0.00699 |
| *Stenotrophomonas maltophilia* HAMBI 2659 | $-3.9\times10^{-5}$ | $7.7\times10^{-5}$ | $1.2\times10^{-4}$ | [-0.001, 0.001] | 0.748 | 0.935 |
| *Niabella yanshanensis* HAMBI 3031 | $2.5\times10^{-5}$ | $1.4\times10^{-4}$ | $1.2\times10^{-4}$ | [$-8.2\times10^{-5}$, $3.7\times10^{-4}$] | 0.467 | 0.7 |
| *Microvirga lotononidis* HAMBI 3237 | $2.3\times10^{-6}$ | 0.000 | $-2.3\times10^{-6}$ | [$-6.8\times10^{-6}$, 0.000] | 1 | 1 |

**Supplementary Table S10. Resistance filtering across streptomycin concentrations.** Within each focal-species history, each non-zero dose was compared with the matched no-antibiotic value using an exact sign-flip test over *N*=8 constructed landscapes. Holm adjustment is across the three non-zero doses for each metric and history. The final two columns give the evolved-minus-ancestral difference at each dose, tested by exact allocation randomization of the 16 constructed landscapes with Holm adjustment across the four doses; these values are identical for the two history rows of a dose because they describe the contrast between them.

| History | Metric | Strep. | Dose − 0 | *P* exact | *P* Holm | *Δ* history | *P* Holm (history) |
|---|---|---|---|---|---|---|---|
| ancestral | MIC-class score | 2 | -0.199 | 0.0078 | 0.0234 | -0.110 | 0.00249 |
| ancestral | MIC-class score | 20 | 1.415 | 0.0078 | 0.0234 | +0.247 | 0.00249 |
| ancestral | MIC-class score | 200 | 1.881 | 0.0078 | 0.0234 | -0.166 | 0.00062 |
| ancestral | High-MIC share | 2 | -0.092 | 0.0078 | 0.0234 | -0.041 | 0.0014 |
| ancestral | High-MIC share | 20 | 0.429 | 0.0078 | 0.0234 | +0.091 | 0.00202 |
| ancestral | High-MIC share | 200 | 0.595 | 0.0078 | 0.0234 | -0.053 | 0.0014 |
| evolved | MIC-class score | 2 | -0.111 | 0.0859 | 0.0859 | -0.110 | 0.00249 |
| evolved | MIC-class score | 20 | 1.860 | 0.0078 | 0.0234 | +0.247 | 0.00249 |
| evolved | MIC-class score | 200 | 1.914 | 0.0078 | 0.0234 | -0.166 | 0.00062 |
| evolved | High-MIC share | 2 | -0.069 | 0.0156 | 0.0234 | -0.041 | 0.0014 |
| evolved | High-MIC share | 20 | 0.583 | 0.0078 | 0.0234 | +0.091 | 0.00202 |
| evolved | High-MIC share | 200 | 0.605 | 0.0078 | 0.0234 | -0.053 | 0.0014 |

**Supplementary Table S11. Sensitivity analysis relating ancestral MIC class to species antibiotic response.** Spearman ρ relates species MIC class to mean ΔCLR response. *P* values are from 200,000 permutations of MIC labels across the nine species of the CLR basis; Holm adjustment is across doses within each history.

| History | Strep. | Spearman ρ | *P* permutation | *P* Holm |
|---|---|---|---|---|
| ancestral | 2 | -0.098 | 0.801 | 0.801 |
| ancestral | 20 | 0.775 | 0.0224 | 0.0671 |
| ancestral | 200 | 0.677 | 0.0556 | 0.111 |
| evolved | 2 | 0.138 | 0.726 | 0.726 |
| evolved | 20 | 0.775 | 0.0237 | 0.0712 |
| evolved | 200 | 0.507 | 0.166 | 0.332 |

**Supplementary Table S12. Primary growth-trait model for competitive position.** Replicate-stratified two-predictor linear model: $q_i$ ~ $z$(relative maximum growth rate) + $z$(relative lag time), fitted independently to $N$=8 constructed landscapes per history over 0, 2 and 20 µg $ml^{-1}$. Values are mean replicate partial slopes with bootstrap 95% CIs. Within-history effects use exact sign-flip tests; history differences use exact allocation randomization. Holm adjustment is across the two traits.

| Trait | Ancestral β [95% CI] | *P* Holm | Evolved β [95% CI] | *P* Holm | Δβ history [95% CI] | *P* Holm |
|---|---|---|---|---|---|---|
| Maximum growth rate | 0.865 [0.840, 0.893] | 0.0156 | 0.717 [0.703, 0.734] | 0.0156 | -0.148 [-0.179, -0.118] | 0.000311 |
| Lag time | -0.399 [-0.441, -0.353] | 0.0156 | -0.547 [-0.593, -0.475] | 0.0156 | -0.148 [-0.218, -0.065] | 0.00637 |

**Supplementary Table S13. Dose-resolved growth-trait models for competitive position.** The same replicate-stratified two-predictor linear model was fitted separately at 0, 2 and 20 µg $ml^{-1}$. Values are mean partial slopes with bootstrap 95% CIs. Exact sign-flip tests assess within-history slopes and exact allocation randomization tests history differences; Holm adjustment is across the two traits within each dose.

| Strep. | Trait | Ancestral β [95% CI] | *P* Holm | Evolved β [95% CI] | *P* Holm | Δβ history [95% CI] | *P* Holm |
|---|---|---|---|---|---|---|---|
| 0 | Maximum growth rate | 1.027 [1.003, 1.053] | 0.0156 | 1.253 [1.231, 1.275] | 0.0156 | 0.226 [0.192, 0.259] | 0.000311 |
| 0 | Lag time | -0.736 [-0.789, -0.647] | 0.0156 | -0.813 [-0.868, -0.730] | 0.0156 | -0.077 [-0.186, 0.030] | 0.258 |
| 2 | Maximum growth rate | 1.128 [1.091, 1.161] | 0.0156 | 1.158 [1.125, 1.194] | 0.0156 | 0.030 [-0.017, 0.080] | 0.276 |
| 2 | Lag time | -0.191 [-0.234, -0.152] | 0.0156 | -0.580 [-0.640, -0.477] | 0.0156 | -0.389 [-0.470, -0.274] | 0.000932 |
| 20 | Maximum growth rate | 0.445 [0.399, 0.505] | 0.0156 | -0.077 [-0.118, -0.037] | 0.0156 | -0.522 [-0.594, -0.459] | 0.000311 |
| 20 | Lag time | -0.052 [-0.126, 0.064] | 0.383 | -0.181 [-0.220, -0.131] | 0.0156 | -0.129 [-0.255, -0.035] | 0.0177 |

**Supplementary Table S14. Dose-resolved variance partition for competitive position.** Within each constructed landscape and dose, joint $R^2$ from the rate-plus-lag model was partitioned by the two-predictor Shapley/LMG decomposition. Values are means across *N*=8 constructed landscapes. Percentages in parentheses are each trait's share of the explained variance; this table is descriptive and does not introduce additional hypothesis tests.

| History | Strep. | Joint $R^2$ | Growth-rate contribution | Lag-time contribution |
|---|---|---|---|---|
| ancestral | 0 | 0.243 | 0.148 (61.1%) | 0.095 (38.9%) |
| ancestral | 2 | 0.139 | 0.136 (97.9%) | 0.0030 (2.1%) |
| ancestral | 20 | 0.021 | 0.018 (91.5%) | 0.0023 (8.5%) |
| evolved | 0 | 0.454 | 0.361 (79.5%) | 0.094 (20.5%) |
| evolved | 2 | 0.498 | 0.322 (65.7%) | 0.176 (34.3%) |
| evolved | 20 | 0.016 | 0.0045 (28.7%) | 0.011 (71.3%) |
